# A circadian transcriptional sub-network and *EARLY FLOWERING 3* control timing of senescence and grain nutrition in bread wheat

**DOI:** 10.1101/2024.02.19.580927

**Authors:** Christopher R. Buckley, Joshua M. Boyte, Robert L. Albiston, Jessica Hyles, Jesse T Beasley, Alexander AT Johnson, Ben Trevaskis, Alexandre Fournier-Level, Michael J. Haydon

**Affiliations:** School of BioSciences, University of Melbourne, Parkville, VIC 3010, Australia; CSIRO Agriculture and Food, GPO Box 1700, Canberra, ACT 2601, Australia

**Keywords:** *Triticum aestivum*, circadian clock, leaf senescence, GPC, chronotype, grain nutrition

## Abstract

Circadian clocks control daily and seasonal timing of physiology and development. Because of their influence on photoperiodic flowering, variants in circadian clock genes have been selected for phenology during domestication of cereal crops. To explore the potential impact of this genetic variation on circadian-regulated traits, we investigated the relationship of the circadian clock and leaf senescence in hexaploid bread wheat. Phenotyping of a collection of elite wheat cultivars identified significant variation in circadian rhythms which was associated with timing of senescence and nutrient mobilisation efficiency. RNA sequencing revealed substantial reorganisation of the circadian-regulated transcriptome during senescence and a transcriptional sub-network representing a link between the circadian oscillator and regulators of leaf senescence. We used genotypes of multiple circadian clock genes to assign cultivars to ‘chronotypes’, which could be used to predict circadian-regulated phenotypes. This identified a deletion variant of *EARLY FLOWERING 3-D1 (ELF3-D1)* attributed to a phenology locus, *Earliness per se (Eps-D1),* and we used near-isogenic lines (NILs) to show that it affects timing of senescence and grain protein content (GPC). Thus, there are potential consequences of circadian clock genes selected for phenology on other valuable crop traits.

## Introduction

Leaf senescence is a controlled process involving the degradation of chlorophyll, downregulation of photosynthesis and the transport of remobilised nutrients to nascent tissues (Gan and Amasino, 1997). Efficient mobilisation of nutrients from vegetative to reproductive tissues during leaf senescence is a key determinant of plant fitness. The relative timing of senescence affects a trade-off between resource accumulation and usage (Distelfeld et al., 2014) and can be adjusted to balance these processes for adaptation to different environments (Estiarte and Peñuelas, 2015; Chen et al., 2020; Bucher and Römermann, 2021). In cereal crops, the timing of senescence has a major impact on yield traits. In bread wheat (*Triticum aestivum*), high grain protein content (GPC) is a valuable trait which can be achieved with earlier senescence, but typically comes at a cost of lower yield (Simmonds, 1995). Furthermore, these relationships can be environment-dependent (Christopher et al., 2016; Christopher et al., 2018; Alhabbar et al., 2018). Increasing nutrient content without yield loss across multiple environments is a priority target for crop breeding.

The circadian clock is a biological timing mechanism, comprised of a highly influential gene regulatory network that functions to synchronise development and physiology with daily and seasonal environmental cycles. Plant circadian clocks strongly affect metabolism, transport and signalling underlying growth and development (Sanchez and Kay, 2016). In numerous cereal crops, the role of the circadian clock in photoperiodic flowering has led to widespread selection of circadian clock gene variants (Bendix et al., 2015), although the impact of these variants on other circadian-controlled traits has not been explored.

Since the circadian clock influences timing of senescence in *Arabidopsis thaliana* (Sakuraba et al., 2014; Zhang et al., 2018; Kim et al., 2018), we sought to explore the role of the circadian clock in wheat in controlling leaf senescence and determining grain nutrient content. We compiled a panel of 25 wheat cultivars and found these traits covaried with phenotypic and genotypic variation in the circadian clock. We compared the circadian transcriptomes of mature and senescent flag leaves and used genotypic information for circadian clock genes to define chronotypes within the wheat panel. These suggested a role for the clock gene *EARLY FLOWERING 3-D1* (*ELF3-D1*) and using near isogenic lines (NILs) of a widespread deletion variant in *ELF3-D1,* we found a functional contribution to timing of senescence and GPC.

## Material and Methods

### Plant materials

Phenotyping experiments in year one and two used a panel of 15 elite Australian wheat cultivars (Calingiri, Cobra, Corack, Gladius, Hydra, Mace, Magenta, Scout, Sunguard, Suntop, Supreme, Viking, Wyalkatchem, Yitpi and Zen) which were obtained from Australian Grains Genebank (AGG). In the third and fourth years, a 25-cultivar circadian diversity panel was developed by including cultivars from the OzWheat diversity panel developed by the Commonwealth Scientific and Industrial Research Organisation (CSIRO). These additional cultivars were Axe, Bayonet, Buckley, Cascades, EGA Wedgetail, Insignia, Lillimur, Takari, Wilgoyne and Yarralinka.

Four pairs of NILs with variation at *TaELF3-D1* were derived from a large NIL population with phenology gene variation developed by CSIRO (Boden et al., 2015; Steinfort et al., 2017). The NILs were generated by crossing donors of phenology genes with the cultivar Sunstate, which carries the *TaELF3-D1del* allele. This was followed by five rounds of recurrent backcrossing to obtain average 97% genetic identity between NILs.

### Growth conditions

For glasshouse experiments, seeds were sown in July-August (Parkville, Victoria). Growth room and growth cabinet experiments were set to 12 h light (200-280 µmol m^-2^ s^-1^, 20°C), 12 h dark (15°C). In all phenotyping experiments, seeds were chilled at 4°C for 48 h and sown in a mix of sterilised potting soil with 4.5 g/L Osmocote 6-9 month slow-release fertiliser. Full details of growth conditions can be found in Supplemental Methods.

### Delayed leaf fluorescence (DF) imaging

Unless specified otherwise, DF was imaged in the second leaves of plants at the 5-leaf stage. In all experiments, 2.5 cm sections were dissected from excised leaves (cut 3 cm from the leaf tip) and placed on 12 cm^2^ 0.5% agar (type M; Sigma-Aldrich) Petri plates, arranged according to a randomised controlled block design. Leaf sections were imaged using a Retiga LUMO charge-coupled device (CCD) camera (Teledyne Photometrics) fitted with a 25 mm f/0.95 lens housed within light-tight dark boxes. Light (>45 µmol m^-2^s^-1^) was provided by two LB3 red/blue LED panels (Photek). The lights and camera were controlled by beanshell (bsh) scripts in µManager (version 2.0 (Edelstein et al., 2014)). DF was measured in 120 h of continuous light (LL) conditions by 1 min camera exposure in the dark, following 59 min of light. Camera properties were binning = 4x4, gain = 1, readout rate = 0.650195 MHz 16 bit. DF was measured in leaf sections from image stacks as integrated density using ImageJ (NIH). Timeseries data were uploaded to BioDare2 (Zielinski et al., 2014) for period analysis of background-subtracted data using the fast Fourier transform non-linear least squares (FFT*-*NLLS) method (Plautz et al., 1997). Data were detrended according to the amplitude and baseline detrending method and analysis was performed using a 24 - 96 h window. Estimates of period that fell outside of the range of 24 h ± 5 h or had relative amplitude error (RAE) > 0.6 were excluded. Time series intensity data presented are Z-score normalised.

### Quantifying time to senescence

Leaf chlorophyll content was recorded using a handheld SPAD meter (Konica Minolta). Measurements were taken three times/week from heading (full emergence of ear). In year two, five SPAD values per plant were measured from the midpoint of the flag leaf of the main tiller and the median value was recorded. In year four, ten SPAD values per plant were measured across the whole flag leaf of the main tiller and the mean value was recorded. Time to senescence was defined as the number of days after anthesis (DAA) to reach 50% of the maximum chlorophyll content. Plants with time to senescence values three standard deviations above or below the overall mean were excluded.

### Nutrient profiling of grain and leaf samples

Whole flag leaves were harvested at anthesis from six plants of each genotype and mature grains were harvested from another six plants. Flag leaf harvest was performed between 9 am and 12 pm (approximately 2-5 h after dawn). For grain samples, all ripened grain heads from each plant were harvested. Thousand grain weight (TGW) was recorded by counting and weighing 50 seeds from each sample. Leaf and grain samples were dried at 65°C for 7 days. Threshed grain samples were sieved with a 2 mm slotted sieve and then milled with a Tube Mill ‘Control’ (Ika). Each sample was ground for 90 seconds at 25,000 rpm. Dried leaf samples were milled using a TissueLyser (Qiagen), with each sample milled for 2 min at 21.5 Hz with a single 18 mm stainless steel ball bearing. Powdered leaf and grain samples were digested in HNO_3_ and ion concentration was determined by inductively coupled plasma optical emission spectroscopy (ICP-OES). Total remobilisation efficiency was calculated as the mean normalised ratio of seed-to-leaf concentration across eight elements (P, S, K, Ca, Mg, Fe, Zn, Mn). Total protein of grain samples was extracted following the protocol of Dupont et al. (2011) and quantified by bicinchoninic acid (BCA) assay.

### Defining chronotypes

Clock gene markers were identified by searching for SNPs in the OzWheat diversity panel (Hyles, 2021). Genotyping of the entire OzWheat panel has been performed with allele-specific markers for agronomically important genes for phenology (e.g. *Ppd-1* and *ELF3*) and around 26K SNPs were identified by transcriptome sequencing (crown tissue from seedlings) of the panel. The OzWheat polymorphism database was filtered against a set of putative wheat clock genes (Supplemental Table 1), and novel non-synonymous SNPs in clock genes were retained. Kompetitive allele specific PCR (KASP) genotyping of clock gene markers (*TaPpd-B1, TaPpd-D1, TaELF3-D1*) and novel SNP markers (*TaPRR73-A1, TaPRR59-B1, TaTOC1-B1, TaLUX-B1*) was completed for the circadian diversity panel. Homoeologue-specific primers for each clock gene marker were designed using PolyMarker (Ramirez-Gonzalez et al., 2015) (Supplemental Table 6).

Genotype data for the seven clock gene markers from cultivars in the OzWheat panel plus additional cultivars in this study were used as input for *k*-modes categorical clustering, using only cultivars with complete genotypic data for the seven genes. To choose the value for *k*, the total within-cluster simple-matching distance was calculated and plotted for a range of possible values of *k* (2–10). *k*-modes clustering was performed using the R/klaR package (Weihs et al., 2005). Chronotypes were visualised in space by multiple correspondence analysis (MCA), performed using the R/FactoMineR package (Lê et al., 2008). To estimate the number of dimensions to retain for MCA, *k*-fold cross-validation was performed using the R/missMDA package (Josse and Husson, 2016), and the number of dimensions (3) that produced the smallest mean square error of prediction (MSEP) was used for MCA.

### RNA-sequencing

Wheat seeds of the Australian cultivar Mace were sown in a single growth room in two staggered batches, 28 days apart. Leaves were sampled concurrently when ‘mature’ plants were at ear emergence (Zadok’s stage 51-59) and ‘senescent’ plants had evident chlorophyll degradation in flag leaves. At dawn (*Zeitgeber* Time 0; ZT0), the growth room was set to continuous light and temperature (20°C) and whole flag leaves from the primary tiller of four plants were collected every 2 h from ZT45 to ZT91 (48 h). Samples were immediately flash frozen in liquid nitrogen and stored at -80°C.

Frozen leaf tissue was ground to a fine powder in liquid nitrogen-cooled mortar and pestles. Lysate buffer was added to the tissue (350 µL per 100 mg of tissue) and the mixture was vortexed immediately. RNA was extracted using ISOLATE II RNA Plant Kit (Meridian Bioscience). RNA from two individual leaf samples was equally pooled to provide two replicates for sequencing. Library preparation and RNA-sequencing was performed by a commercial provider (Azenta). Library preparations were constructed using NEBNext Ultra RNA Library Prep Kit for Illumina (New England Biolabs; NEB). Libraries with different indices were multiplexed and sequenced on an Illumina HiSeq platform (Illumina, San Diego, CA). Sequencing was carried out using a 2x150-bp paired-end (PE) configuration. An average of 24 million 150-bp PE reads were generated for each library.

Reads were filtered for quality and mapped to the *T. aestivum* cv. Mace (PGSBv2.1) genome sequence and annotation using STAR with default parameters (Dobin et al., 2013). Mapped reads were quantified and normalised using the cuffquant and cuffnorm modules of Cufflinks (Trapnell et al., 2010). Normalised counts (fragments per kilobase transcript per million reads; FPKM) outputted from cuffnorm were used for downstream analyses. Only transcripts that were expressed (i.e. FPKM > 0) at least once in each 24 h period (ZT45-ZT67 and ZT69-ZT91) were retained.

### Parameterisation of transcript circadian rhythms

The packages BIO_CYCLE (Agostinelli et al., 2016) and r/MetaCycle (Wu et al., 2016) were used to estimate rhythmicity and parameterise circadian rhythms of transcripts. BIO_CYCLE was run using default parameters, with minimum and maximum expected period values of 20 and 28 h. MetaCycle incorporates different methods for rhythmicity detection to generate a combined prediction of rhythmicity. JTK_CYCLE and Lomb-Scargle (LS) were selected for integration by MetaCycle. Default settings were maintained for all other parameters. Transcripts with *q* < 0.05 were retained from both BIO_CYCLE and MetaCycle, and the intersection of rhythmic genes from these datasets was used for downstream analyses. All circadian rhythm parameter estimates (period, phase, amplitude, periodicity) presented are derived from BIO_CYCLE. Phase values are adjusted to period (phase = phase(24/period)). The mature-to-senescent change in each of these parameters at the gene level (Δperiod, Δphase and Δamplitude) are used for several analyses. ΔPeriod and Δphase are calculated as the difference between the mature and senescent value of these parameters for a given gene (e.g. Mature period - senescent period). ΔAmplitude is the log ratio of mature and senescent amplitude for a given gene (i.e. log(Mature amplitude: senescent amplitude)).

### Gene ontology (GO) term and TF family enrichment analysis

GO terms for the *T. aestivum* cv. Mace (PGSBv2.1) annotation were sourced from Ensembl Plants (Cunningham et al., 2022). r/topGO was used to perform GO enrichment analysis of various gene sets using the ‘classic’ algorithm (Alexa et al., 2006). Fisher’s exact test was selected to compare the enrichment of GO terms in each gene set to a gene universe of 87,493 unique wheat genes with GO term annotations. For enrichment analyses of TF families, Chinese Spring wheat genes annotated as TFs were sourced from https://opendata.earlham.ac.uk/wheat/under_license/toronto/Ramirez-Gonzalez_etal_2018-06025-Transcriptome-Landscape/data/data_tables/ (Ramírez-González et al., 2018). Mace orthologues of each gene were assigned through orthology data available from Ensembl Plants (Cunningham et al., 2022). For enrichment analyses, 2 x 2 contingency tables were built for each TF family and analysed with Fisher’s exact test (from r/stats). Enrichment analysis of TF families that are only rhythmic in senescent tissue was performed against all rhythmic TFs (rhythmic in senescent, mature or both). Enrichment analysis of differentially rhythmic TFs was performed against all rhythmic TFs (rhythmic in both tissues).

### Cis-regulatory element enrichment

Promoter sequences (1000 bp upstream of the start codon) were obtained from all wheat genes using the ‘retrieve sequences’ tool from RSAT (Regulatory Sequence Analysis Tools) (Nguyen et al., 2018). Promoter sequences were subsetted into groups of interest (e.g. short period genes), and these groups were used as input for *cis*-regulatory element enrichment analysis. For this, promoter sequences from the target set of genes were used as input for RSAT peak-motifs oligo-analysis (Thomas-Chollier et al., 2012), with all genes that were rhythmic in either mature or senescent flag leaves used as the control set. Peak-motifs settings were kept as default, and significantly enriched oligos with E-value < 0.05 were retained. For each significantly enriched oligo, motif matching was performed with STAMP (Similarity, Tree-building, and Alignment of Motif Profiles) (Mahony and Benos, 2007) against the PLACE database of plant motifs (Higo et al., 1998), using default settings. The top match for each oligo (with smallest *p*-value) was retained.

### Differential expression and differential rhythmicity

LimoRhyde (linear models for rhythmicity, design) was used to test for statistically significant changes in rhythmicity between mature and senescent tissue (differential rhythmicity; DR) (Singer and Hughey, 2019). LimoRhyde first decomposes *Zeitgeber* Time into sine and cosine with a period of 24 h. Then, transcripts that exhibit DR are identified using a linear model that tests for significant interactions between developmental stage and the time components. LimoRhyde was also used to identify differentially expressed (DE) transcripts. For this, a linear model tests for interactions between developmental stage without interaction of the time components(Singer and Hughey, 2019). Code used to perform LimoRhyde’s functions was developed from a vignette from the LimoRhyde GitHub repository (https://github.com/hugheylab/limorhyde). *p*-values were adjusted by the Benjamini-Hochberg procedure with a false discovery rate FDR = 5% (Benjamini and Hochberg, 1995). DR and DE genes were assigned using a *q*-value threshold of 0.05. DE genes have log2(fold change) > 1.2 or < -1.2. To allow comparison of circadian rhythm parameters (i.e. period, phase, amplitude), the DR gene set was filtered against the set of genes that are rhythmic in both tissues.

### STRING protein-protein network analysis

Protein-protein interactions of DR wheat TFs were analysed using STRING (Szklarczyk et al., 2023). STRING uses an early genome assembly and annotation for wheat (CSS; International Wheat Genome Sequencing Consortium (IWGSC), 2014). CSS gene IDs for the DR gene list were acquired via BLAST searches within STRING, though not all Mace gene annotations had an equivalent locus in the CSS assembly. DR TFs with STRING entries were used to build a protein-protein network in STRING. The network was built using both physical and functional protein interactions, and the minimum weighted interaction score was set to 0.4.

### Clustering of oscillator transcript responses

Mace orthologues of putative wheat circadian oscillator genes from Chinese Spring were identified using orthology data from Ensembl Plants (Cunningham et al., 2022) (Supplemental Table 2). The list of genes was filtered to include only transcripts that are rhythmic in both mature and senescent tissue. Principal component analysis (PCA) was performed for the 59 clock genes in this list using three variables: mature-to-senescent changes in circadian period, phase, and amplitude (Δperiod, Δphase and Δamplitude). The three PCs from this analysis were used as input for *k*-means clustering. To determine an optimal value of *k*, the ratio of within-cluster sum-of-squares (WCSS) to total sum-of-squares (TSS) was plotted against a range of potential *k* values (1-8). PCA and *k*-means were performed using r/stats.

## Results

### Circadian period and timing of senescence covary in Australian wheat cultivars

Since the circadian clock influences leaf senescence in Arabidopsis (Sakuraba et al., 2014; Zhang et al., 2018; Kim et al., 2018) and the pace of circadian rhythms shortens with age (Kim et al., 2016; Rees et al., 2019), we investigated the relationship between phenotypic variation of circadian rhythms in wheat and the timing of senescence. Using delayed leaf fluorescence (DF) (Figure 1A), we first measured circadian rhythms in the second leaf of three wheat cultivars 14 days after sowing (DAS; seedling), 33 DAS (5-leaf stage) and 44 DAS (end of tillering). Relative amplitude error (RAE; a proxy for rhythm robustness) did not significantly differ by leaf age (Figure 1B), but circadian period was significantly shorter in leaves of older plants (Figure 1C). This demonstrates that changes occur in the circadian clock of wheat plants during ageing.

**Figure 1:**
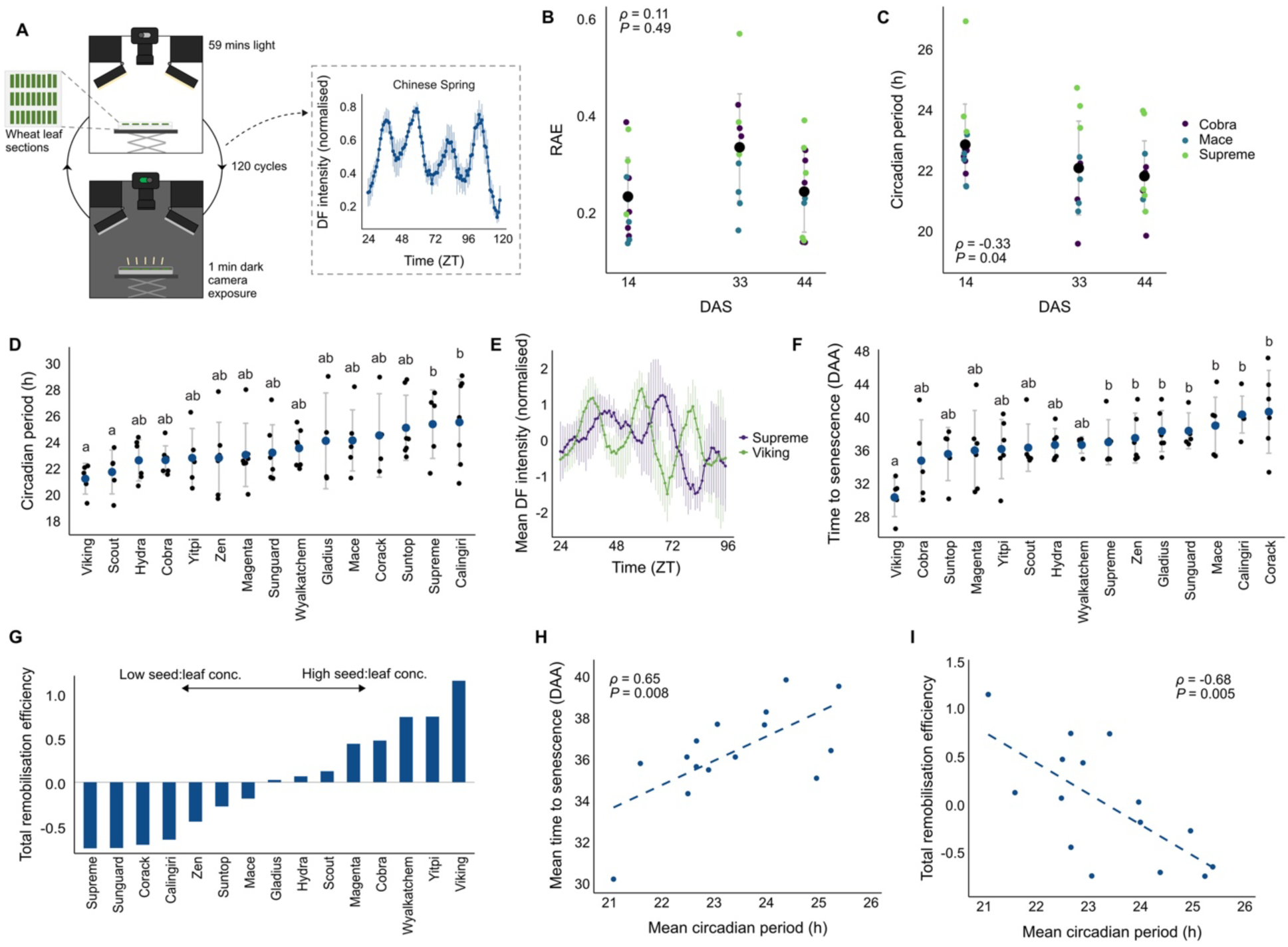
Circadian period of wheat cultivars is associated with senescence and grain nutrition. **A,** Overview of delayed fluorescence (DF) workflow for measuring wheat circadian rhythms. **B,** Relative amplitude error (RAE) and **C**, circadian period of DF rhythms in 2^nd^ leaf of three cultivars at 14, 33 and 44 days after sowing (DAS). Significance of relationship with age (DAS) assessed by Pearson’s product-moment correlation. Error bars represent ±sd, n=15. **D,** Circadian period of DF in 15 Australian wheat cultivars in year one. Letters indicate significant differences as determined by Welch’s *t*-test, *P* < 0.05. Error bars represent ±sd, n=8. **E,** Circadian rhythms of DF in the Supreme and Viking. Error bars represent ±sd, n=8. **F,** Time to senescence, (days after anthesis; DAA) in 15 wheat cultivars. Letters indicate significant differences as determined by one-way ANOVA followed by Tukey’s HSD; *P* < 0.05. Error bars represent ±sd, n=6. **G,** Total nutrient remobilisation efficiency (ratio of leaf-to-seed nutrient content, see methods) in 15 wheat cultivars. **H**, Mean circadian period (year one) versus mean time to senescence and **I,** total remobilisation efficiency in 15 wheat cultivars. Significance assessed by Pearson’s product-moment correlation.

To assess the extent of phenotypic variation of circadian rhythms in wheat, we chose 15 elite cultivars grown across diverse environments of Australia (Supplemental Figure 1) and measured circadian rhythms of DF in glasshouse-grown plants in two years. Significant differences in mean circadian period were detected between cultivars in each year (Figure 1D, Supplemental Figure 2A, Supplemental Data 1). In year 1, mean circadian period differed by over four hours over the panel (Figure 1D,E). Circadian period was mostly consistent within cultivars over both years, except for Viking and Scout, which might suggest strong gene-by-environment interactions in these cultivars (Supplemental Figure 2B). Nevertheless, there is robust variation in circadian rhythms within elite wheat germplasm which could influence circadian-regulated agronomic traits.

To measure the timing of senescence in the 15 wheat cultivars, we monitored the decline of chlorophyll content in flag leaves after anthesis (Supplemental Data 1). There were significant differences in the timing of senescence between cultivars (Figure 1F), with a maximum mean difference of 10.3 days. We also measured total ion content of mature flag leaves and harvested grain to calculate nutrient mobilisation efficiency (Supplemental Data 1). We detected substantial variation in mobilisation efficiency between cultivars (Figure 1G; Supplemental Figure 3) and observed a strongly negative correlation with timing of senescence (Supplemental Figure 4). Timing of senescence and nutrient mobilisation efficiency were both significantly correlated with circadian period in year one (Figure 1H,I), such that cultivars with shorter period tended to senesce earlier and had more efficient nutrient mobilisation. The relationship of these traits to circadian period was weaker in year two due to a strong year-effect for Scout and Viking. Without these cultivars, a stronger correlation was detected (Supplemental Figure 2C,D). We also measured total GPC, an important trait which depends on nutrient mobilisation (Supplemental Figure 5A, Supplemental Data 1). GPC was significantly negatively correlated with circadian period in year two, but not year one (Supplemental Figure 5B,C). Overall, our data suggest strong relationships between circadian period, leaf ageing and senescence, nutrient mobilisation efficiency and GPC in elite wheat cultivars.

### Reshaping of the circadian transcriptome during leaf senescence

To explore transcriptional changes in the circadian network during leaf senescence we compared the transcriptomes of mature and senescent flag leaves in *cv.* Mace over 48 h in continuous light (Figure 2A). Principal component analysis (PCA) separated each individual transcriptome by developmental stage (Supplemental Figure 6A) and time of day (Supplemental Figure 6B-E). We used BIO_CYCLE (Agostinelli et al., 2016) and MetaCycle (Wu et al., 2016) to detect rhythmic transcripts and estimated their period, phase and amplitude (Supplemental Data 2). A total of 11,705 expressed transcripts were called rhythmic in mature flag leaves (15.7% of 74550 expressed) by both algorithms and 16,906 transcripts were rhythmic in senescent flag leaves (22.8% of 74272 expressed) (Figure 2B). Since fewer rhythmic genes were detected in mature flag leaves compared to senescent flag leaves, the changes in the circadian transcriptome during senescence are unlikely to be due to a general decay in gene expression. We performed gene ontology (GO) enrichment analysis of transcripts that are rhythmic only in senescent flag leaves (Supplemental Data 3). Significantly enriched terms included “protein transport”, “peptide metabolic process” and “nitrogen compound transport” (Figure 2C), suggesting that the circadian network is connected to the reprogramming of metabolism and transport during senescence.

**Figure 2:**
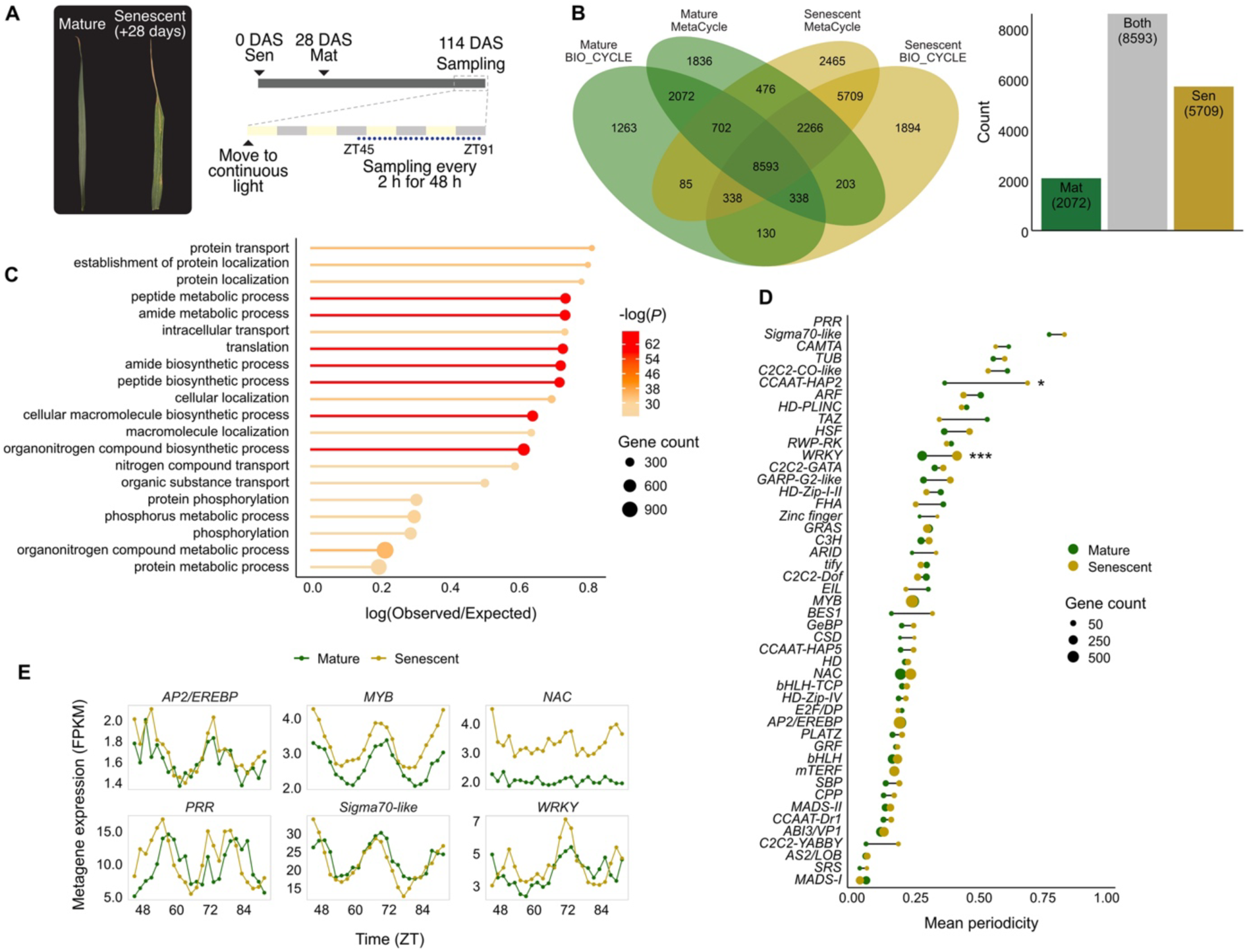
Circadian transcriptional network increases its output at leaf senescence. **A,** Sampling design of comparative circadian senescence transcriptome. ‘Sen’ and ‘Mat’ refer to the sowing dates of plants that were harvested for senescent and mature flag leaves, respectively. **B,** Number of transcripts determined to be rhythmic by BIO_CYCLE (Agostinelli et al., 2016) and MetaCycle (Wu et al., 2016). **C,** Gene ontology term enrichment analysis of transcripts that are uniquely rhythmic in senescent flag leaves. **D,** Comparison of mean periodicity of transcription factor families determined by BIO_CYCLE (0 = arrhythmic). Asterisks indicate significant differences as determined by Welch’s *t*-test, * = *P* < 0.05, *** = *P* < 0.001. **E,** Metagene expression (average across entire gene family) of selected transcription factor families.

To identify potential regulators of the circadian transcriptome during leaf senescence, we compared periodicity (BIO_CYCLE’s measure of rhythmicity, from 0 to 1) across families of transcription factors (TFs) (Ramírez-González et al., 2018). *PSEUDO RESPONSE REGULATORs* (*PRRs*), which are core circadian oscillator genes (Nakamichi et al., 2005), were the most rhythmic family in both tissue types, followed by Sigma70-like TFs, which regulate rhythmic chloroplast gene expression in Arabidopsis (Figure 2D) (Noordally et al., 2013). Mean periodicity of the *WRKY* and *CCAAT-HAP2* families was significantly higher in senescent tissue than mature tissue (Figure 2D). To summarise the overall changes in TF gene expression, we calculated metagene expression values of entire TF gene families (Figure 2E) and observed a clear increase of *WRKY* metagene rhythmicity during senescence. By contrast, a similar change in rhythmicity was not observed for other TF families with roles in senescence (e.g. *NAC, AP2* and *MYB* (Cao et al., 2023)), although we did detect a phase advance of metagenes for the *MYB, PRR* and *Sigma70* families (Figure 2E). *WRKY* genes are known as regulators of leaf senescence (Guo and Gan, 2005) but our data suggests the influence of the circadian clock on their regulation might have been underestimated.

Among the 8,593 rhythmic transcripts in both tissues, the mean circadian period was ∼0.5 h shorter in senescent flag leaves compared to mature flag leaves (Figure 3A; Supplemental Figure 7B). This is consistent with the expectation for shorter period in older leaves (Figure 1C). In addition, a circadian phase advance of ∼0.5 h (corrected for period) was also detected in senescent leaves (Supplemental Figure 7A,B). Amplitude differences were small between the two developmental stages, although mean amplitude was higher in mature tissue (Supplemental Figure 7C).

**Figure 3:**
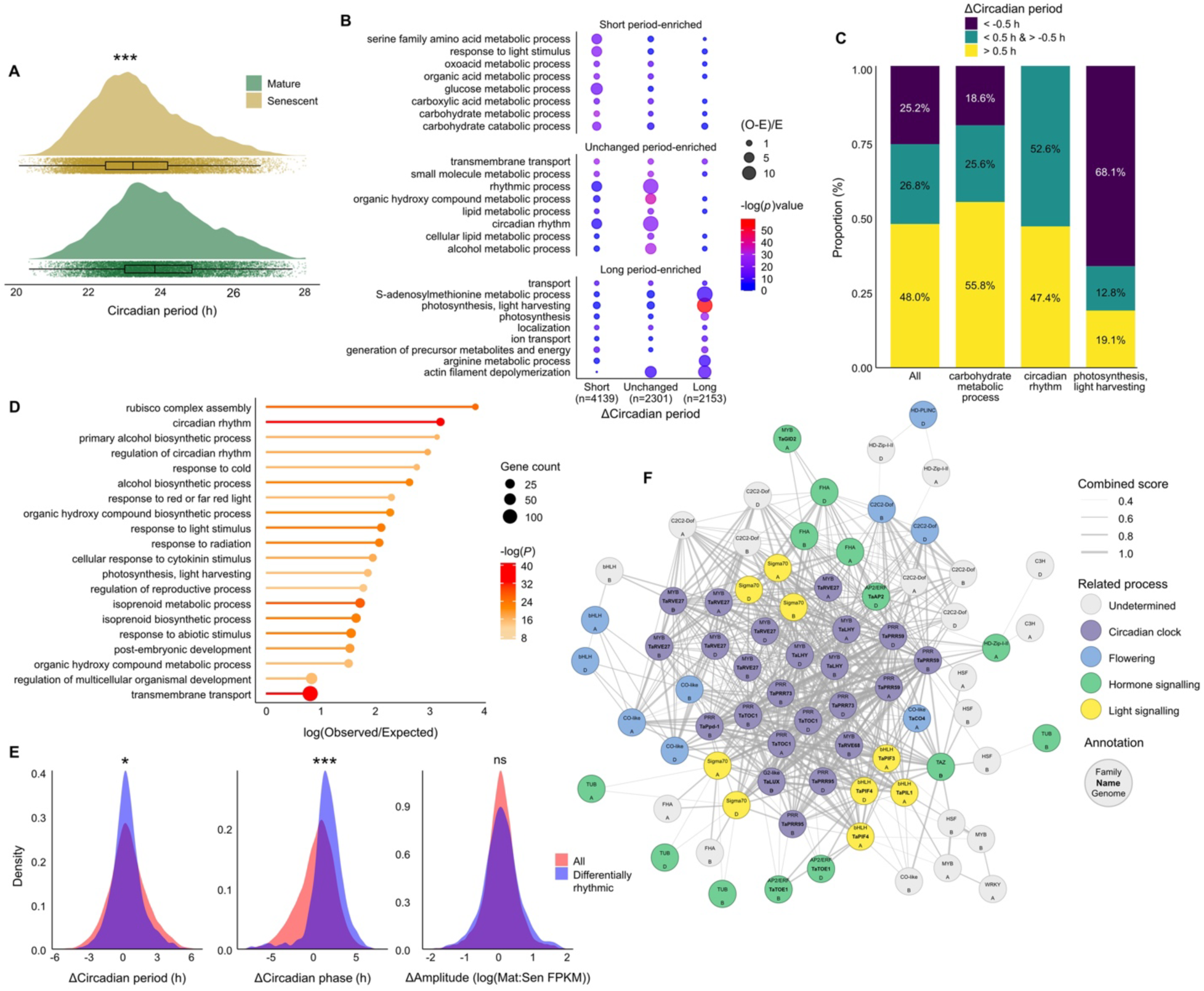
A circadian-regulated transcriptional subnetwork in senescent leaves. **A,** Distribution of circadian period values of all rhythmic transcripts in mature and senescent flag leaves. Asterisks indicate a significant difference between the mean circadian period values, Welch’s *t*-test, *P* < 0.001. **B,** GO term enrichment analysis of transcripts categorised according to the change in circadian period between developmental stages; short period (Δperiod < -0.5 h), long period (Δperiod > 0.5 h), unchanged period (Δperiod < ±0.5 h). O (observed) and E (expected) number of transcripts. **C,** Proportions of transcripts in each Δperiod category for a selection of GO terms. **D,** GO term enrichment analysis of differentially rhythmic (DR) genes. **E,** Distribution of changes in circadian rhythm parameters for DR genes. Asterisks indicate significant differences between DR genes and all other rhythmic genes as determined by Welch’s *t*-test, * = *P* < 0.05, *** = *P* < 0.001, ns = not significant. **F,** Putative protein-protein interaction network of DR transcription factors. Combined score reflects the probability of interaction determined by STRING (Szklarczyk et al., 2023). Related processes are manually assigned after literature and orthology database searches.

We noted that there was substantial variation in the period differences for individual transcripts outside the global trend (Figure 3A). To identify patterns in differential period, we performed GO term enrichment of transcripts with shorter (< -0.5 h), longer (> 0.5 h) or unchanged (< ±0.5 h) circadian period in senescent leaves (Supplemental Data 3). Rhythmic genes with shorter period were significantly enriched for GO categories related to primary metabolism (Figure 3B,C; Supplemental Figure 8). Among transcripts with longer period in senescent tissue, terms related to photosynthesis were overrepresented, perhaps reflecting the breakdown of normal rhythmic regulation of photosynthesis (Figure 3B,C; Supplemental Figure 8). The two most strongly enriched GO terms within transcripts with unchanged period were “rhythmic process” and “circadian rhythm” (Figure 3B,C; Supplemental Figure 8). This suggests that the global shortening of period in the senescent transcriptome could be driven by amplitude or phase changes in the circadian oscillator.

To identify other transcriptional changes between mature and senescent leaves, we used LimoRhyde (Singer and Hughey, 2019) to identify differentially expressed (DE) and differentially rhythmic (DR) genes (Supplemental Figure 9, Supplemental Data 3). Among DE genes in senescent leaves, 6,310 upregulated genes were enriched for GO terms for transport and catabolism, and 8,785 downregulated genes were enriched for GO terms related to biosynthesis (Supplemental Figure 10). There were 1,743 DR genes, enriched for GO terms related to photosynthesis, light signalling and circadian rhythms (Figure 3D). When we compared DR genes to the remainder of the rhythmic transcriptome, we observed an advanced mean phase in senescent tissue, rather than change in period or amplitude (Figure 3E).

### A circadian-regulated sub-network in senescent leaves

We hypothesised that the phase-advanced DR genes, which are enriched for the “circadian rhythm” GO term, represent a sub-network driving shorter period of the wider transcriptome. We identified 106 TFs among the DR genes (Supplemental Data 3), which were characterised by large phase advances (Supplemental Figure 11A). Several families of TFs were significantly over-represented, including C2C2-Dof, TAZ, FHA, PRR and Sigma70 (Supplemental Figure 11B). To analyse the interactions between DR TFs, we modelled the putative interactions between members of this gene set using STRING (Szklarczyk et al., 2023). We identified 308 putative functional and physical interactions, which is 7x more than expected (Hypergeo. test, *P* < 1x10^-16^). The network includes genes related to hormone signalling, light signalling and photoperiodic flowering, and is centred around core clock genes; *PRRs*, *REVEILLEs* (*RVEs*), *LUX ARRHYTHMO* (*LUX*) and *LATE ELONGATED HYPOCTYL* (*LHY*) (Figure 3F). Evening elements (EEs), which are *cis*-regulatory motifs recognised by clock TFs (Harmer et al., 2000), were highly significantly enriched among all DR gene promoters (Supplemental Table 1). These results suggest that there is a circadian-controlled transcriptional subnetwork with advanced phase in senescent tissue.

We performed a systematic analysis of circadian oscillator gene expression and identified 59 out of 79 putative circadian oscillator genes (Supplemental Table 2) that were rhythmic in both tissues (Supplemental Figure 12). Mean period difference of this group was 0.52 h, which was similar to the global trend, and suggested that maintenance of period is not uniform across the oscillator (Figure 3B). Strikingly, the mean phase advance of oscillator genes was 1.37 h, which is significantly larger than the global average (Welch’s *t*-test, t (59.5) = 4.21, *P* < 0.001). To categorise the oscillator genes by rhythmic expression patterns, we performed PCA of changes in period, phase and amplitude (Figure 4A). The three PCs were grouped into six transcript clusters of 1-20 genes by *k*-means (Figure 4C). Clusters 5 and 6 are mainly comprised of genes expressed around dawn, including *RVEs*, *NIGHT LIGHT–INDUCIBLE AND CLOCK-REGULATED* (*LNKs*) and *LHY* genes (Figure 4C) and have shorter period in senescing tissue, consistent with the global trend (Figure 4B,D). Clusters 2, 3 and 4 have advanced phase in senescing tissue (Figure 4B), similar to the identified subnetwork (Figure 3F). These are mostly afternoon- or evening-phased genes and include the majority of *PRR* and *LUX* genes (Figure 4C). Amplitude was increased in cluster 2 genes, and period was unchanged in clusters 2 and 4 (Figure 4B,D). Thus, there appear to be distinct changes in oscillator genes during senescence according to their phase of expression and this suggests a restructuring of the circadian oscillator during leaf senescence.

**Figure 4:**
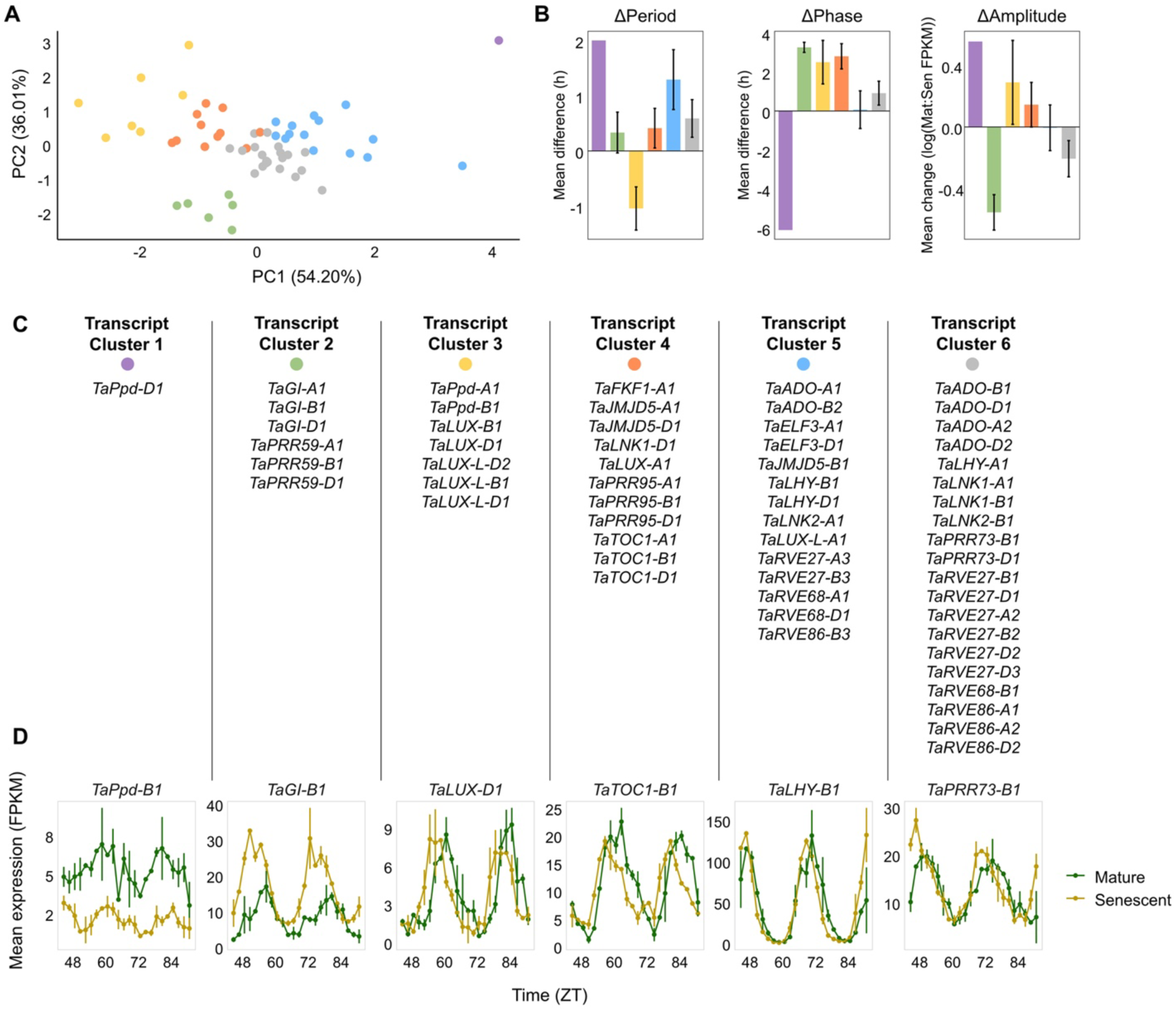
Reshaping of the circadian oscillator during leaf senescence. **A,** PCA of changes in circadian period, phase and amplitude between mature and senescent leaves for putative circadian oscillator genes. PCs generated were used to group transcripts into 6 clusters (represented by different colours). **B,** Mean change in circadian period, phase and amplitude for each transcript cluster. **C,** Names of genes within each transcript cluster. **D,** Transcript circadian rhythms for a representative gene from each cluster. Error bars represent ±sd.

### Identification of chronotypes within elite wheat cultivars

Since the circadian clock appears to strongly influence the wheat transcriptome during leaf senescence, we sought to maximise circadian gene variation within the wheat panel for further phenotyping. We explored SNP data for the OzWheat diversity panel comprised of 283 Australian cultivars. We found four novel non-synonymous SNPs in conserved regions of clock genes from distinct transcript clusters (Figure 3C); *TaPRR59-B1* (Cluster 2), *TaLUX-B1* (Cluster 3), *TaTOC1-B1* (Cluster 4) and *TaPRR73-A1* (Cluster 6). We used these SNPs and three major polymorphic clock gene loci in wheat (*TaPpd-B1*, *TaPpd-D1* and *TaELF3-D1* (Beales et al., 2007; Cane et al., 2013; Wittern et al., 2023)), to categorise cultivars into six chronotypes (Figure 5A,B; Supplemental Table 3) and compiled a circadian diversity panel of 25 cultivars, representing all six chronotypes (Supplemental Table 4), which were phenotyped for circadian period and timing of senescence (Figure 5E,F; Supplemental Figure 13, Supplemental Data 1). There were significant differences between chronotypes for both circadian period and time to senescence (Figure 5C,D). In turn, grain filling was also affected, as thousand grain weight (TGW) differed significantly between chronotypes (Supplemental Figure 14A,B, Supplemental Data 1). However, there were no significant differences between chronotypes for GPC (Supplemental Figure 14C,D, Supplemental Data 1). Although the circadian period values do not correlate with time to senescence and TGW for all chronotypes, chronotype 5 exhibited short period, early senescence and low TGW, while chronotype 6 showed both longer period and later senescence (Figure 5C,D; Supplemental Figure 14).

**Figure 5:**
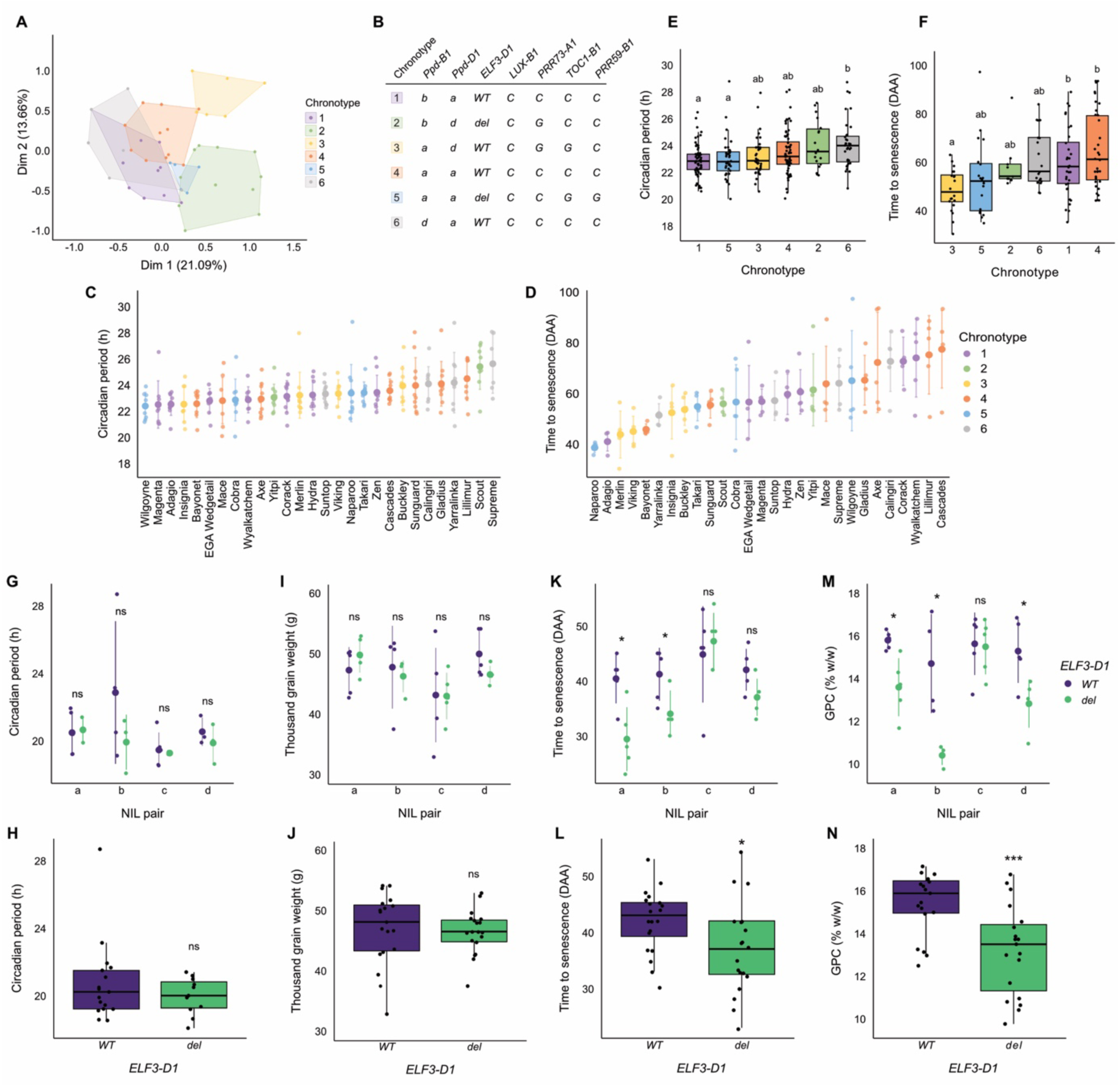
Wheat clock gene variation affects time to senescence and GPC. **A,** Cultivars from the OzWheat diversity panel were grouped into chronotypes by *k*-modes clustering based on their genotype at seven clock gene markers. Multiple correspondence analysis was used to visualise clusters in space. **B,** Mode allelic variation of each cluster (see Supplemental Table 3 for explanation of alleles). **C,** Circadian period (n=10) and **D**, time to senescence (n=5) of 25 wheat cultivars in a circadian diversity panel. Error bars represent ±sd. **E,** Circadian period and **F,** time to senescence for the 25 cultivars by chronotype defined in **A**. Letters indicate significant differences as determined by one-way ANOVA followed by Tukey’s HSD; *P* < 0.05. **G,H,** Circadian period, **I,J,** thousand grain weight (TGW), **K,L,** time to senescence and **M,N,** grain protein content (GPC) of near isogenic lines (NILs) for variation at *TaELF3-D1*. Individual NIL pairs are shown in **G,I,K,M.** Error bars represent ±sd, n=5 . Values for all four NILs with functional (*WT*) and *ELF3-D1del (del)* are combined in **H,J,L,N.** Asterisks indicate significant differences between *WT* and *del* alleles as determined by Welch’s *t*-test, * = *P* < 0.05, *** = *P* < 0.001, ns = not significant.

### *ELF3-D1* deletion impacts timing of senescence and GPC

Chronotypes 2 and 5 both have a deletion in *TaELF3-D1* and relatively early senescence. To test the potential for this circadian polymorphism on timing of leaf senescence, we used four pairs of near-isogenic lines (NILs) that differ by variation at *TaELF3-D1* (Supplemental Table 3,4, Supplemental Data 1). Across the four pairs of NILs circadian period was not significantly different (Figure 5G,H) and neither was TGW (Figure 5I,J). However, early senescence was detected for the *TaELF3-D1* deletion in three out of four NIL pairs (Figure 5K,L). Similarly, GPC was reduced by the *TaELF3-D1* deletion in the same three NIL pairs (Figure 5M,N). The exception is NIL pair c, which carries a different genotype for *TaPpd-D1* (Supplemental Table 5), a putative downstream target of *ELF3*. Thus, the absence of an effect in this NIL pair could be explained by an epistatic interaction.

## Discussion

We have used a diverse collection of elite wheat cultivars and comparative transcriptomics to explore the relationship between circadian clock function and the process of leaf senescence in wheat and assess its impact on grain quality. Our data reveal significant phenotypic variation in circadian rhythms within wheat cultivars and identify strong correlations with timing of leaf senescence, nutrient mobilisation efficiency and GPC. Comparison of the circadian transcriptome in mature and senescent leaves identified a circadian-controlled subnetwork, centred around several core clock genes and connected to possible transcriptional regulators of senescence. Using genotypic information for circadian clock genes in these wheat cultivars we were able to define chronotypes which we show could potentially be used to predict circadian phenotypes and yield traits. Phenotyping of NILs for an extant deletion variant in *TaELF3-D1* revealed a functional contribution of this core circadian clock gene to timing of senescence and GPC.

ELF3 is a core component of the circadian oscillator as part of the evening complex (EC), a transcriptional repressor complex also comprised of ELF4 and LUX in Arabidopsis (Nusinow et al., 2011). Null mutants in *ELF3* in Arabidopsis and double null mutants in tetraploid wheat have arrhythmic circadian phenotypes (Hicks et al., 1996; Wittern et al., 2023). Hypomorphic alleles of *elf3* in Arabidopsis have short period phenotypes (Kolmos et al., 2011; Anwer et al., 2014). Chronotypes 2 and 5 have a *TaELF3-D1* deletion, and chronotype 5 had a relatively short period (Figure 5). Since we did not detect significant differences between the *TaELF3-D1* NIL pairs, it seems unlikely that this variant alone causes the short period phenotype. Notwithstanding that the *TaELF3-D1* deletion might be recessive to functional A and B homoeologues for circadian period, there are several genotypic differences in clock genes between these two chronotypes, so the short period of chronotype 5 could be due to genetic interactions between different clock alleles. For example, chronotypes 2 and 5 have different alleles at *TaPpd-D1* and *TaPpd-B1*, which are putative regulatory targets of the EC (Alvarez et al., 2023).

We found a significant effect of the *TaELF3-D1* deletion on timing of leaf senescence and GPC for three out of four NIL pairs (Figure 5). This suggests that the widespread *TaELF3-D1* deletion, attributed to *Earliness per se (Eps-D1)* locus (Zikhali et al., 2016; Wittern et al., 2023), has a potential negative effect on grain quality, independent of any effect on circadian period. The exception was NIL pair c, which has a promoter deletion within *TaPpd-D1,* in contrast to the functional alleles present in the other three (Supplemental Table 5). Thus, the effect of *TaELF3-D1* on senescence might depend on the genotype at *TaPpd-1.* Notably, *TaPpd-B1* is among the *PRR* genes identified within the transcriptional subnetwork (Figure 3) which we predict to make a regulatory contribution to leaf senescence.

Increased GPC is commonly associated with reduced yield in cereal crops (Simmonds, 1995) and this relationship has been attributed to timing of senescence (Distelfeld et al., 2014). TGW, which is an important yield component, was not significantly different between *TaELF3-D1* NILs (Figure 5), but accurate estimates will require multi-year field trials. These will also be required to better assess the robustness of the relationship between circadian period and timing of senescence, since we detected substantial gene-by-environment effects for some cultivars (Supplemental Figure 2).

The overall trend in the rhythmic transcriptome was of shorter circadian period in senescent tissue (Figure 3) which is consistent with expectations of accelerating rhythms with age (Kim et al., 2016; Rees et al., 2019) (Figure 1). However, there was substantial variation across the transcriptome. For example, photosynthesis genes had a longer period, which could be explained by general decay in expression and an uncoupling from the core oscillator. Of particular note, genes that were relatively unchanged for period were enriched for circadian genes and instead are characterised by an advanced phase (Figure 3). Indeed, three of the circadian transcript clusters we defined had advanced phase in senescent tissue, and include numerous *PRR* genes, EC genes and *GIGANTEA* (Figure 4). We propose that the phase adjustment of these oscillator genes could be driving the period change of the wider transcriptome. Interestingly, *ELF3* genes and *TaPpd-D1* are among the few oscillator genes that have a substantially shorter period in senescent leaves and might represent a regulatory link between the adjusted oscillator and senescence-regulated genes.

Our findings have potentially important implications for breeders and growers because they suggest that extant variation in circadian clock genes in crops, which were selected for their effects on phenology, can have a significant impact on other traits such as grain quality. The extent of these effects is likely to depend on genetic interactions between other circadian loci and so identifying chronotypes within wheat germplasm could be a useful tool to predict phenotypic effects. The TFs within the subnetwork are candidates that could be targeted to fine-tune circadian-regulated leaf senescence without affecting the core circadian clock.

## Acknowledgements

We thank Tina Rathjen for her help with genotyping and genetic analysis and Vanessa Melino for sharing seed. Research was funded by the Research Continuity Scheme from the Faculty of Science, University of Melbourne and the JN Peters Bequest to MJH, and an Alfred Nicholas Fellowship, Melbourne Research Scholarship and Faculty of Science Postgraduate Writing-Up Award to CRB. This research was not funded by the Australian Research Council. Generation of NILs at CSIRO was supported by GRDC project CSP00183.

## Author contributions

MJH conceived the study, CRB and MJH designed experiments, CRB, JMB, RLA and MJH performed experiments, CRB, JH, AFL and MJH analysed data, JH, JTB, AATJ, BT and AFL provided expertise and resources, CRB and MJH wrote the manuscript, JH, BT and AFL reviewed and edited the manuscript.

## Data availability

Sequencing data have been deposited in NCBI Gene Expression Omnibus (GSE259431).

