## Supplemental Information for "A circadian transcriptional sub-network and *EARLY FLOWERING 3* control timing of senescence and grain nutrition in bread wheat"

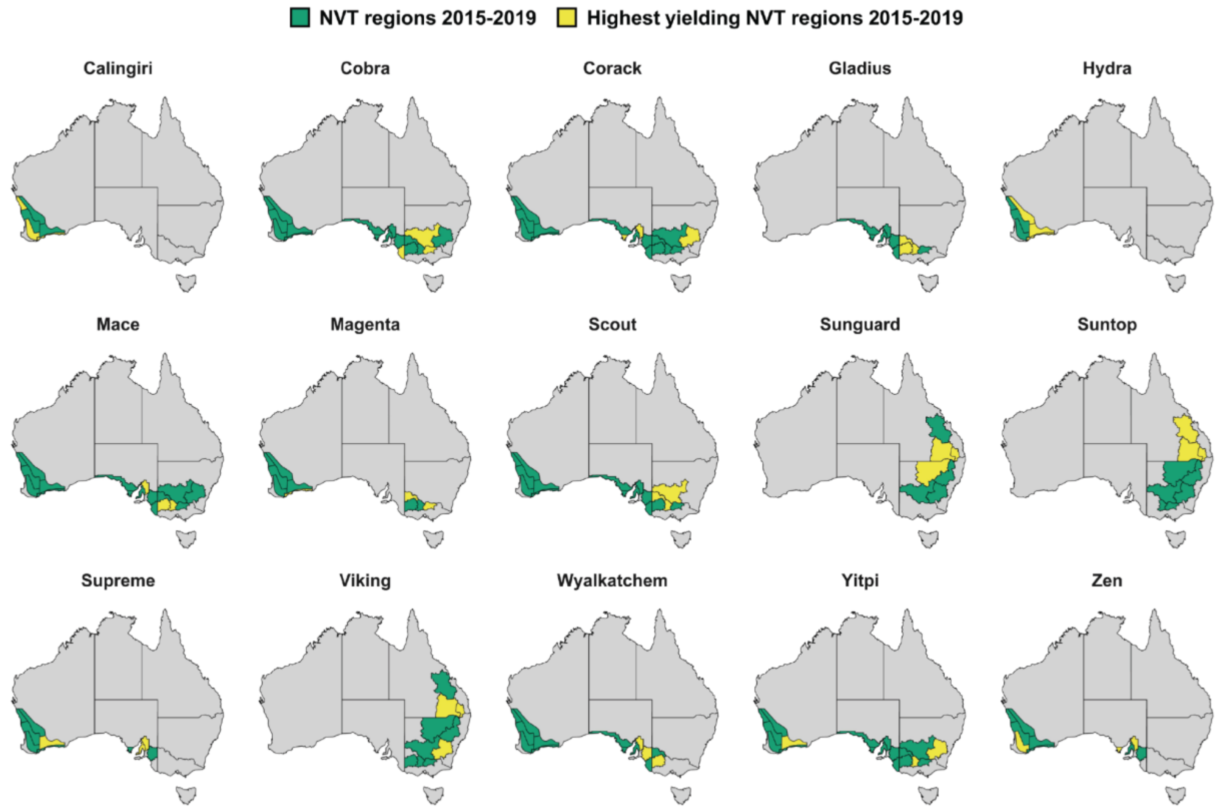


**Supplemental Figure 1: Geographical distribution of 15 elite Australian wheat cultivars.** Growing region data are sourced from Grains Research and Development Corporation (GRDC) National Variety Trials (NVT) main season wheat trial data from 2015-2019 (https://nvt.grdc.com.au/). Regions where cultivars were trialled at least once in the 5-year period are shaded green, whilst the three regions which saw the highest average yield (as a percentage of total average yield across all plots) are shaded yellow. Maps were created using GIS data available from https://www.data.wa.gov.au (for Western Australia Agzones) and https://www.abs.gov.au/ (for Statistical Area Level 2 boundaries, which the other regions are drawn from).

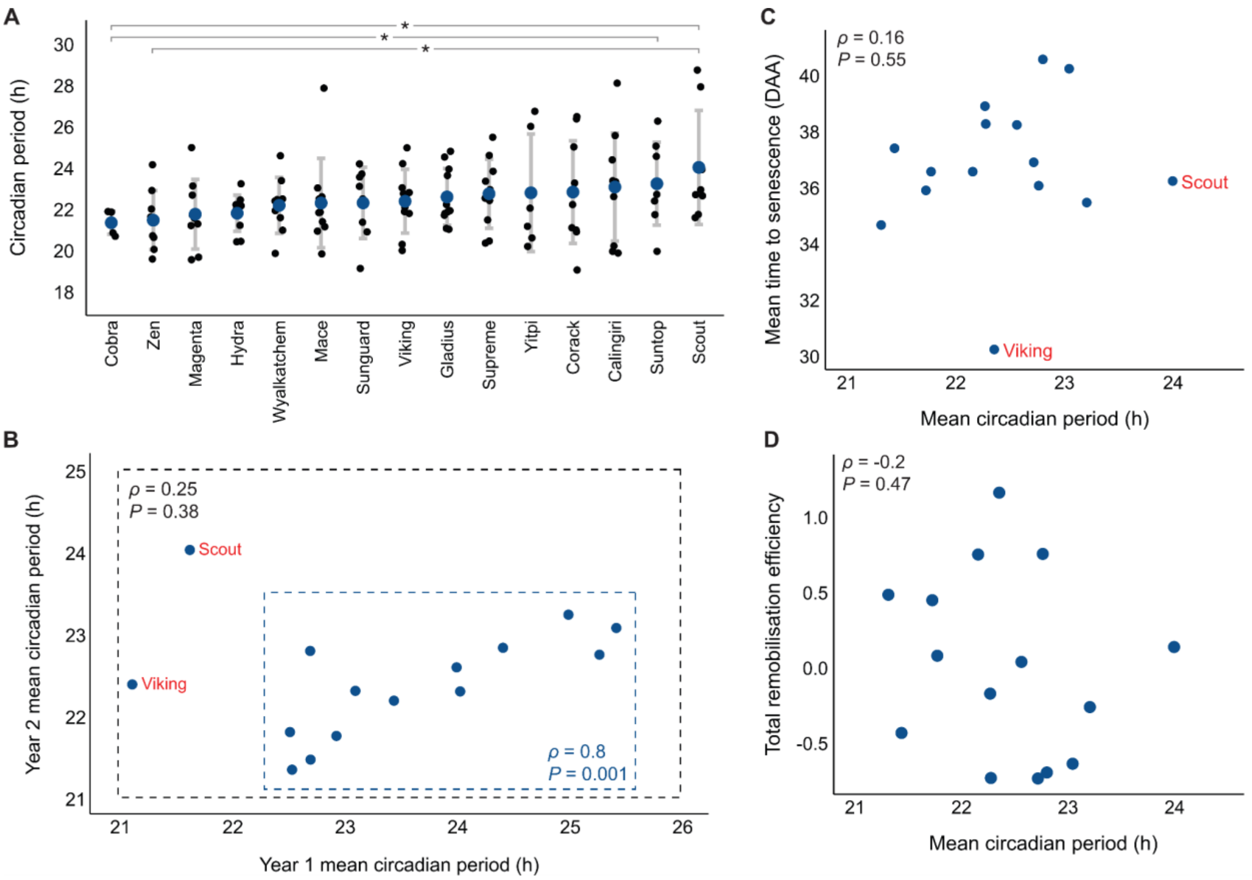


**Supplemental Figure 2: DF circadian period data from year two. A,** Comparison of circadian period from year two across 15 elite Australian wheat cultivars. Letters indicate significant differences as determined by Welch's *t*-test, $P < 0.05$. Error bars represent ±sd, n=12. **B,** Association between circadian period from year one and year two. Analysis of correlation was performed with and without outliers Scout and Viking. **C,** Association of circadian period from year two and time to senescence. **D,** Association of circadian period from year two and total remobilisation efficiency. Significance of association assessed by Pearson's product-moment correlation.

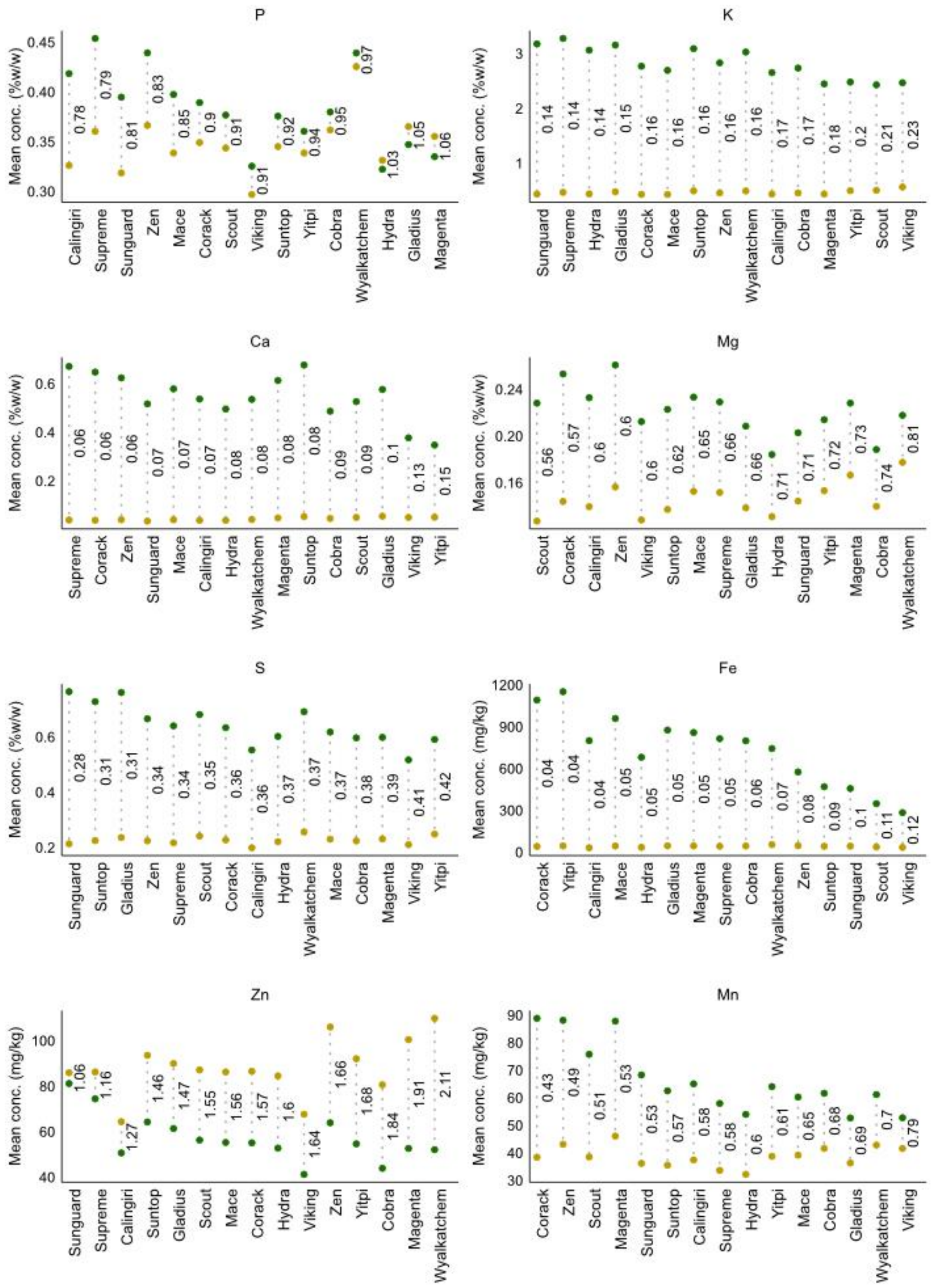


**Supplemental Figure 3: Seed-to-leaf remobilisation efficiency for individual elements.** Mean ion concentration of wheat grain (yellow) and flag leaves (green) for 15 cultivars (n=6). Values next to dashed lines are ratios of seed: leaf concentration.

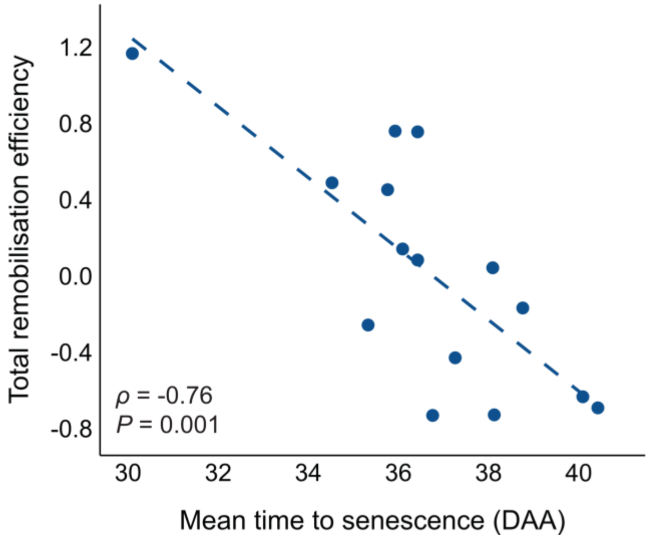


**Supplemental Figure 4: Association between total remobilisation efficiency and time to senescence for 15 cultivars.** Significance assessed by Pearson's product-moment correlation.

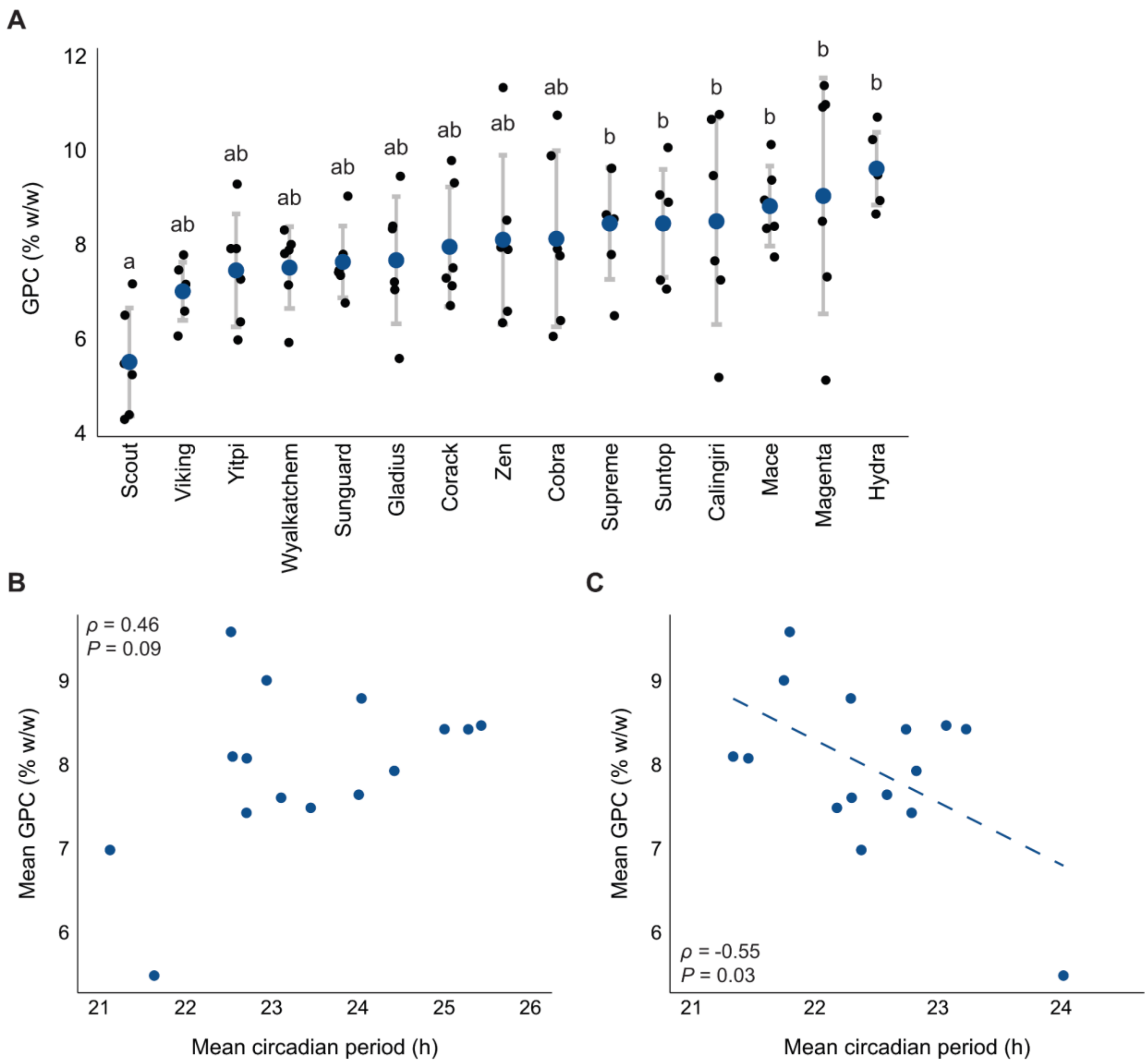


**Supplemental Figure 5: GPC is associated with circadian period. A,** GPC in 15 wheat cultivars. Letters indicate significant differences as determined by one-way ANOVA followed by Tukey's HSD; $P < 0.05$. Error bars represent ± sd, n=6. **B,** Mean circadian period (year 1) versus mean GPC for 15 cultivars. **C,** Mean circadian period (year 2) versus mean GPC in 15 wheat cultivars. Significance of association assessed by Pearson's product-moment correlation.

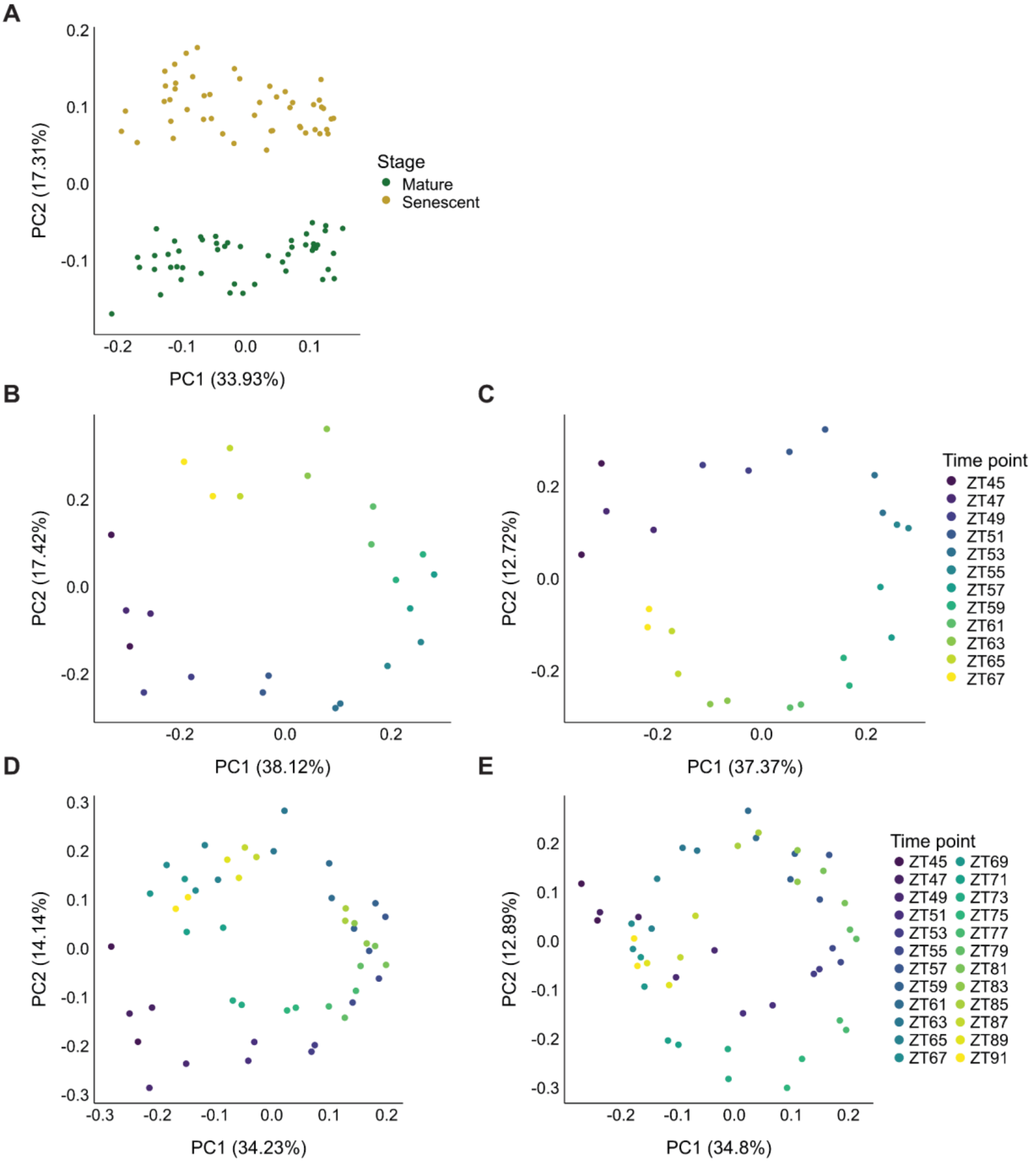


**Supplemental Figure 6: Assessment of transcriptome quality via PCA.** For each PCA, FPKM values from all expressed transcripts in the given dataset were used as input. **A,** Mature versus senescent transcriptomes. **B,** First 24 h (ZT45-ZT67) of mature transcriptomes. **C,** First 24 h (ZT45-ZT67) of senescent transcriptomes. **D,** All mature transcriptomes. **E,** All senescent transcriptomes.

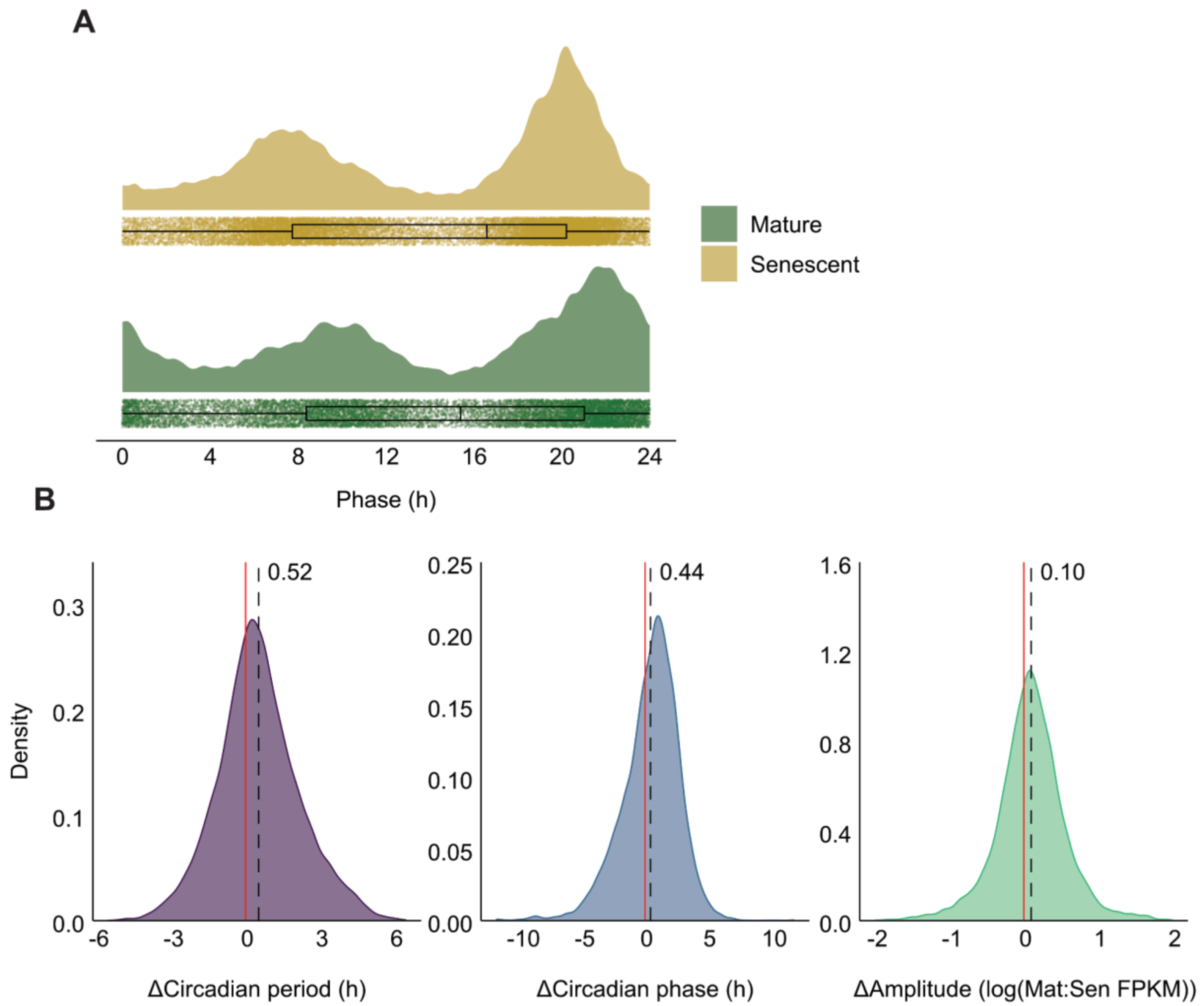


**Supplemental Figure 7: Changes in parameters of circadian rhythms between mature and senescent flag leaves. A,** Distribution of circadian period values of all rhythmic transcripts at each developmental stage. **B,** Distribution of changes in circadian rhythm parameters (circadian period, phase and amplitude) between mature and senescent flag leaves at the gene level. Analysis was performed with 8,593 genes that are rhythmic in both tissues to allow comparison of circadian rhythm parameters for individual genes. Red line indicates 0, dashed line indicates mean.

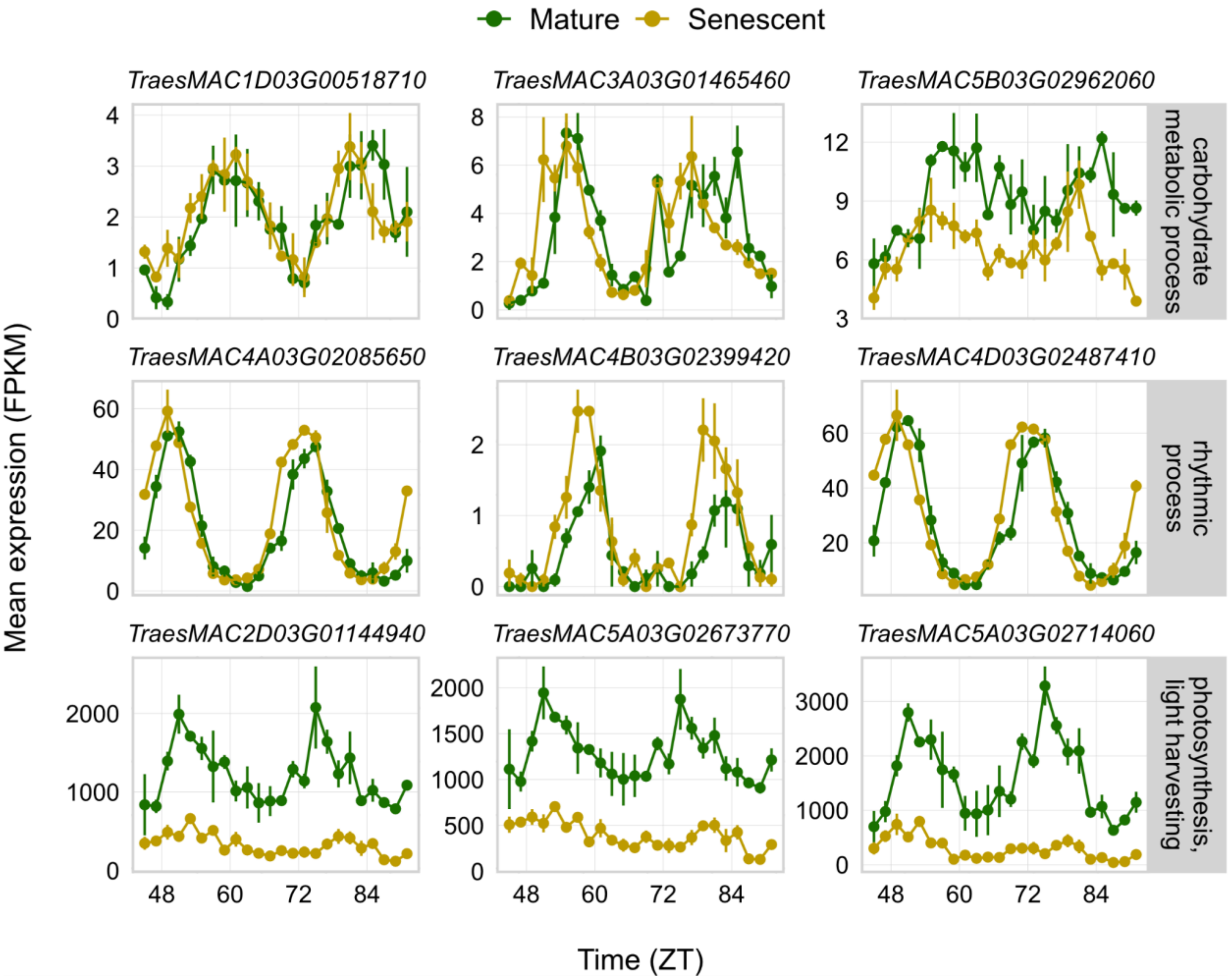


**Supplemental Figure 8: Transcript rhythms for genes within GO terms associated with differential change in circadian period.** GO terms represented correspond to those in Figure 3C. Error bars represent ±sd. Gene annotation or description: 'carbohydrate rhythmic process': *TraesMAC1D03G00518710*, glycosyl hydrolase family protein; *TraesMAC3A03G01465460*, glycosyl hydrolase family protein; *TraesMAC5B03G02962060*, alpha-1,4 glucan phosphorylase. 'Rhythmic process': *TraesMAC4A03G02085650*, *NIGHT LIGHT–INDUCIBLE AND CLOCK-REGULATED 1* (*TaLNK1-A1*); *TraesMAC4B03G02399420*, orthologue of Arabidopsis *COLD-REGULATED GENE 27* (*COR27*); *TraesMAC4D03G02487410*, *TaLNK1-D1*. 'Photosynthesis, light harvesting': *TraesMAC2D03G01144940*, chlorophyll a-b binding protein; *TraesMAC5A03G02673770*, chlorophyll a-b binding protein; *TraesMAC5A03G02714060*, chlorophyll a-b binding protein.

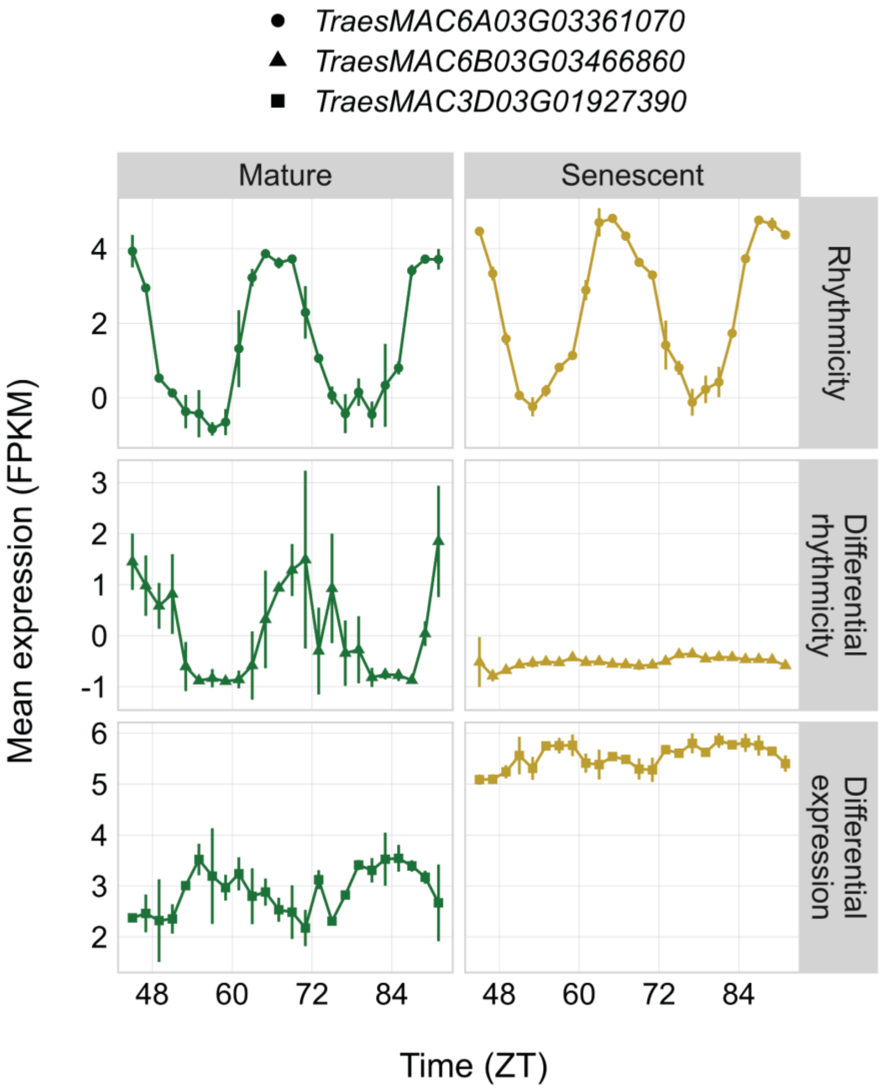


**Supplemental Figure 9: Comparison of the different outputs from LimoRhyde.** The LimoRhyde(Singer and Hughey, 2019) algorithm tests for rhythmicity, differential rhythmicity (DR) and differential expression (DE). Top panel: example expression of a rhythmic gene (note: the rhythmicity feature of LimoRhyde was not used in this study and is included here for illustrative purposes only). Middle panel: example expression of a DR gene. DR genes have statistically significant interactions between the timing of their expression and the developmental stage. Bottom panel: example expression of a DE gene. DE genes have statistically significant interactions between their overall expression and the developmental stage. Error bars represent ±sd.

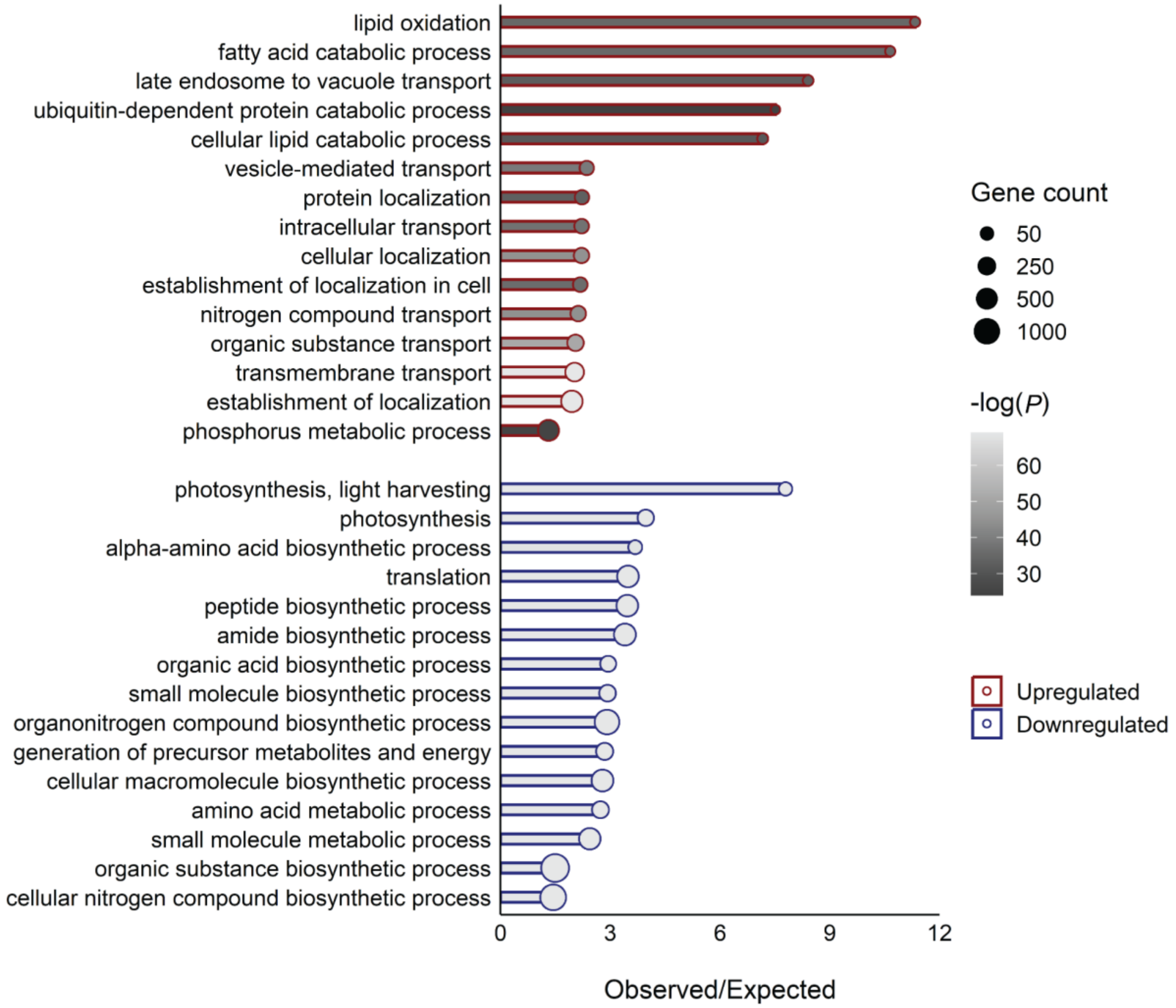


**Supplemental Figure 10: GO term enrichment analysis of differentially expressed genes.** Differentially expressed genes were assigned via LimoRhyde (Singer and Hughey, 2019).

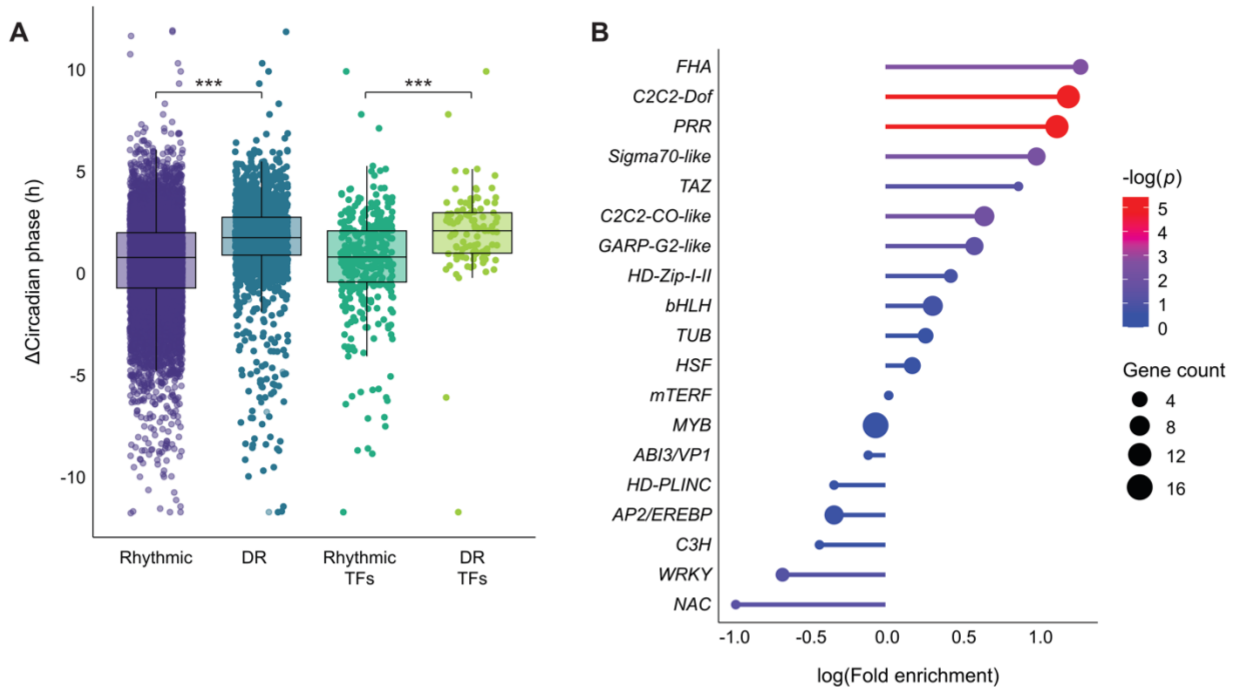


**Supplemental Figure 11: Analysis of differentially rhythmic (DR) transcription factors**. **A,** Change in circadian phase between mature and senescent flag leaves of DR transcription factors (DR TFs) versus all transcripts that are rhythmic at both developmental stages (rhythmic), DR transcripts, and TF transcripts that are rhythmic at both developmental stages (rhythmic TFs). Asterisks indicate significant differences as determined by Welch's *t*-test, $P < 0.001$. **B,** Gene family enrichment analysis of DR TFs.

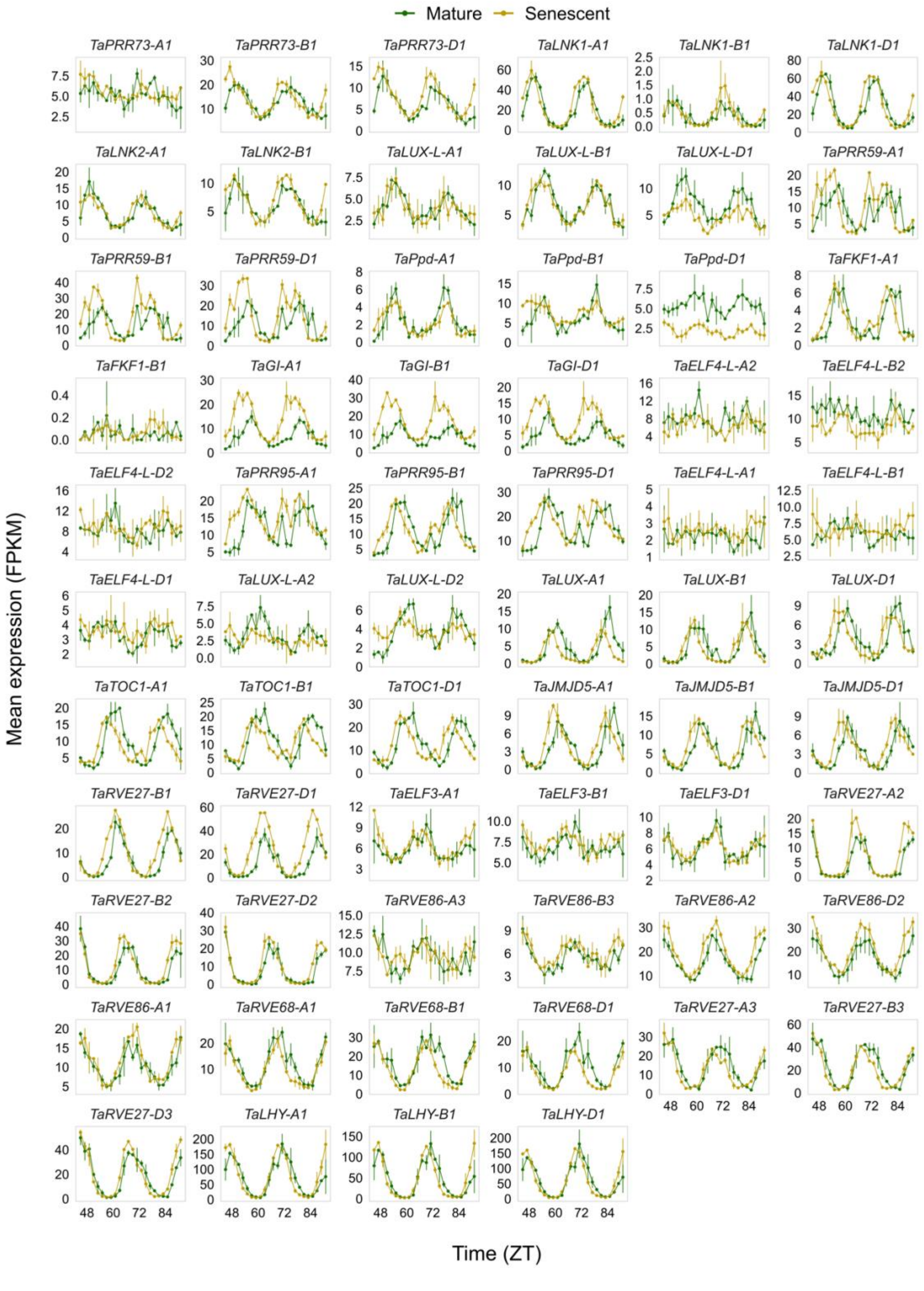


**Supplemental Figure 12: Transcript expression rhythms of clock genes in mature and senescent flag leaves.** Homoeologue triads are ordered by their mean circadian phase across both developmental stages, from ZT0 to ZT24. Error bars represent ± sd.

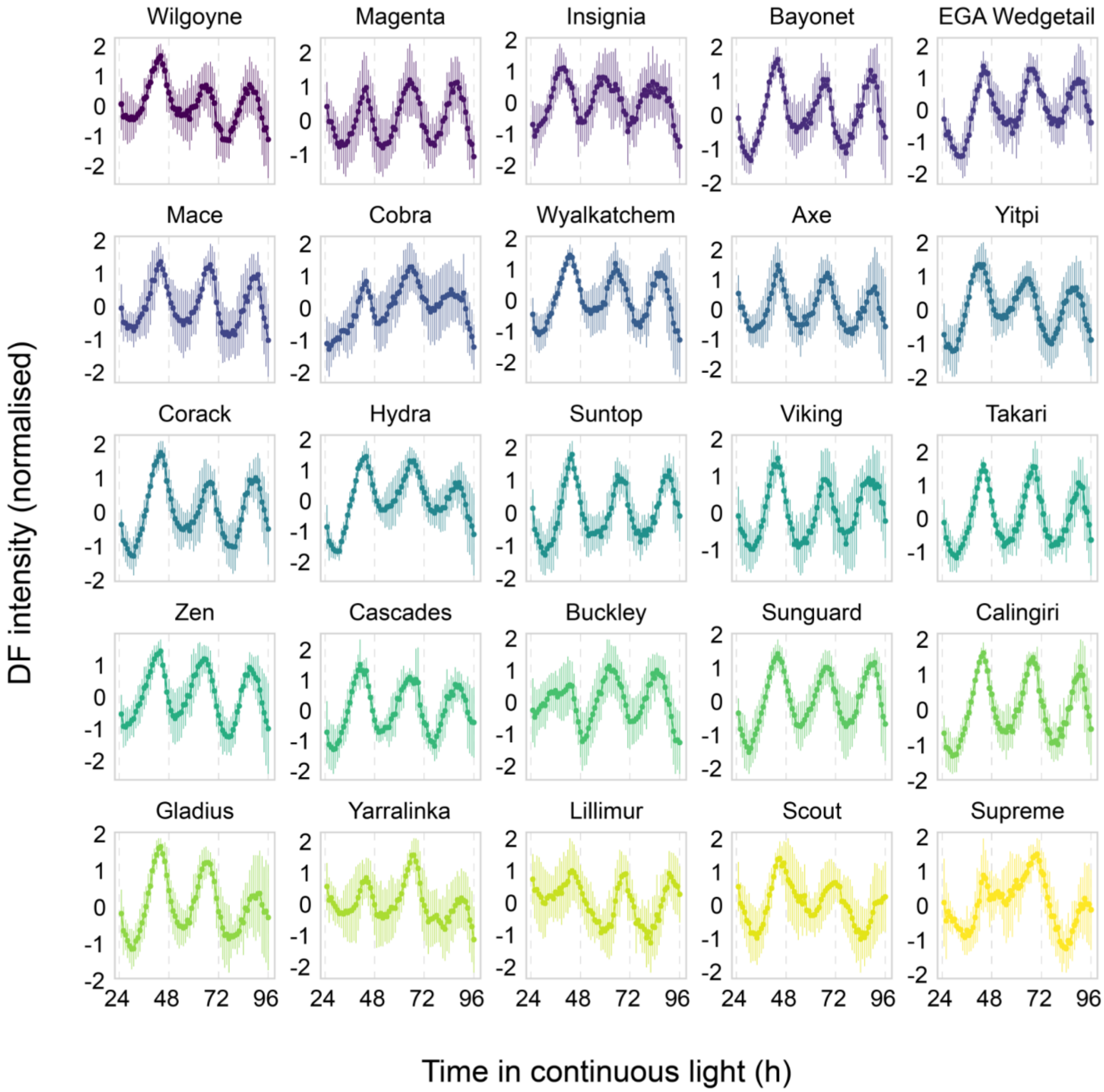


**Supplemental Figure 13: DF circadian rhythms of the circadian diversity panel.** Cultivars are ordered by mean circadian period, from shortest to longest. Error bars represent ±sd, n=10.

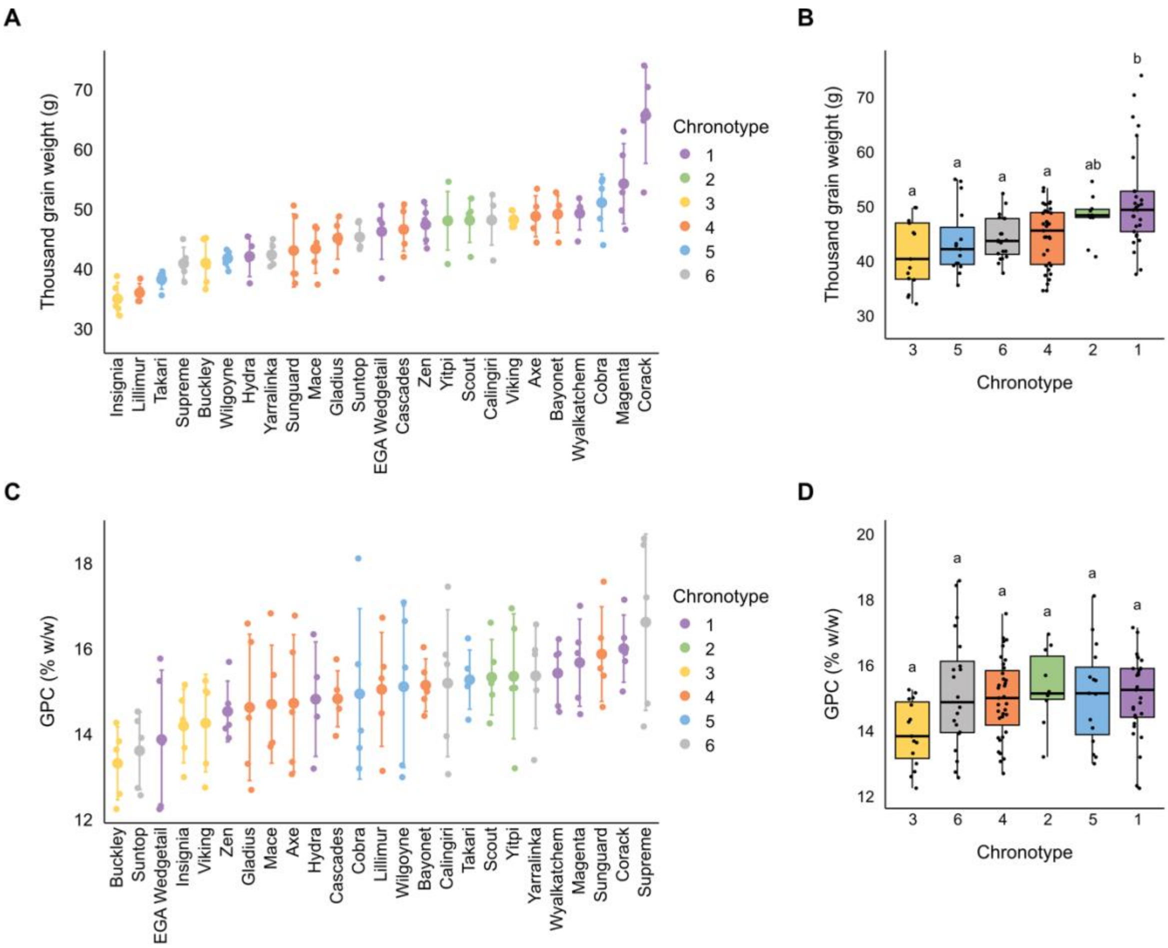


**Supplemental Figure 14: GPC and TGW phenotypes across the circadian diversity panel.** **A,** TGW in 25 wheat cultivars. Error bars are ±sd, n=5. **B,** TGW of 25 cultivars separated by chronotype. **C,** GPC in 25 wheat cultivars. Error bars are ±sd, n=5. **D,** GPC of 25 cultivars separated by chronotype. Letters indicate significant differences as determined by one-way ANOVA followed by Tukey's HSD; $P < 0.05$. Error bars represent ±sd.

**Supplemental Table 1: Overrepresented *cis*-regulatory elements within transcript circadian rhythm gene sets.** For each gene set, promoter sequences (1000 bp upstream of the start codon) were used as input for RSAT (Regulatory Sequence Analysis Tools) peak-motifs oligo-analysis. Each enrichment analysis was performed against a control gene set of all rhythmic genes (rhythmic at either developmental stage). Enriched oligos were matched to known plant cis-regulatory motifs using the motif-matching program STAMP (Similarity, Tree-building, and Alignment of Motif Profiles).

| Gene set | Motif | RSAT sig. * | RSAT E-val † | STAMP top motif match ‡ | STAMP *p*-val § | Motif description | Genes with motif | Max motifs per gene | Gene with max motifs | Wheat gene name or description | Mean motifs per gene |
|---|---|---|---|---|---|---|---|---|---|---|---|
| Mature unique | CGGGCCA | 1.79 | 0.016 | GCBP2ZMGAPC4 | $5.3 \times 10^{-7}$ | Binding site of tobacco nuclear factor (GCBP-2) | 252 (13.35%) | 4 | *TraesMAC7D03G 04403630* | | 1.22 |
| Senescence unique | cCGGCCCA | 2.94 | 0.0011 | PREMOTIFNPCABE | $1.2 \times 10^{-9}$ | Found in Tobacco *CAB* promoter, related to photoregulated expression | 985 (18.77%) | 7 | *TraesMAC1D03G 00452000* | DUF1620-containing WD40-like repeat protein | 1.32 |
| Senescence unique | AGCCCA | 1.69 | 0.02 | SITEIIATCYTC | $3.9 \times 10^{-9}$ | Site II element, found in cytochrome promoters | 1830 (34.87%) | 7 | *TraesMAC2A03G 00810430* | Potassium efflux antiporter | 1.38 |
| Senescence unique | TTCAAA | 1.41 | 0.039 | ERELEE4 | $3.7 \times 10^{-9}$ | Ethylene-responsive element (ERE) of tomato | 2013 (38.36%) | 17 | *TraesMAC2B03G 00909720* | Peptide chain release factor PrfB1 | 1.85 |
| Short period | CCACACG | 2.08 | 0.0083 | ACGTSEED2 | $2.4 \times 10^{-6}$ | "ACGT" motif related to seed germination | 609 (15.52%) | 12 | *TraesMAC4D03G 02443710* | | 1.26 |
| Short period | GCCGCGCCaCG CaCG | 1.46 | 0.035 | ABRE3OSRAB16 | $2.3 \times 10^{-12}$ | ABA-responsive element (ABRE) of rice | 1444 (36.80%) | 11 | *TraesMAC6D03G 03746470* | | 1.99 |
| Short period | CTCCCTC | 1.38 | 0.042 | BOXCPSAS1_3 | $2.4 \times 10^{-6}$ | BOX-C motif from pea asparagine synthetase (*AS1*) gene | 1720 (43.83%) | 36 | *TraesMAC6A03G 03328630* | | 2.26 |
| Unchanged period | GCCACGTGGC | 8.13 | $7.4 \times 10^{-9}$ | ACGTROOT1 | 0 | The 'perfect palindromic' sequence (PA) containing G-box | 492 (22.73%) | 10 | *TraesMAC7D03G 04305940* | | 2.49 |

| | | | | | | | | | | | |
|---|---|---|---|---|---|---|---|---|---|---|---|
| Unchanged period | CGCGGGCC | 1.75 | 0.018 | SORLIP2AT | $1.7x10^{-6}$ | Sequence over-represented in light-induced promoters (SORLIP) | 658 (30.39%) | 9 | *TraesMAC5D03G03158380* | | 1.8 |
| Unchanged period | GCCGaCG | 1.47 | 0.034 | DRECRTCOREAT | $5.4x10^{-8}$ | DRE (dehydration-responsive element) | 663 (30.62%) | 16 | *TraesMAC6A03G03295190* | | 1.8 |
| Unchanged period | CCGGGCC | 1.75 | 0.018 | GCBP2ZMGAPC4 | $5.2x10^{-7}$ | Binding site of tobacco nuclear factor (GCBP-2) | 343 (15.84%) | 10 | *TraesMAC6B03G03571290* | | 1.41 |
| Unchanged period | CGCCACGT | 1.76 | 0.017 | ACGTSEED3 | $6.6x10^{-13}$ | "ACGT" motif related to seed germination | 734 (33.90%) | 5 | *TraesMAC5B03G02931810* | Rhomboid-like protein | 1.45 |
| Unchanged period | CCCGCGC | 1.37 | 0.043 | RE1ASPHYA3 | $2.4x10^{-6}$ | Repressor element associated with oat *phyA3* gene | 437 (20.18%) | 12 | *TraesMAC5B03G02934070* | *TaPRR59-B1* | 1.95 |
| Long period | GGCCCC | 1.96 | 0.011 | AMMORESVDCRNIA1 | $1.2x10^{-8}$ | Motif from Chlamydomonas *Nia1* gene promoter, associated with ammonium response | 616 (30.54%) | 5 | *TraesMAC5D03G03214030* | Non-specific phospholipase C1 | 1.45 |
| DR | AAATATCT | 29.93 | $1.2x10^{-30}$ | EVENINGAT | $1.2x10^{-13}$ | Evening element (EE), binding site of clock genes | 582 (35.25%) | 8 | *TraesMAC7D03G04350940* | *TaJMJD5-D1* | 1.53 |

* RSAT peak-motifs oligo-analysis significance value = log10(E-value)

† RSAT peak-motifs E-value. The *p*-value calculated from the oligo-analysis binomial test, adjusted for expected number of false-positives

‡ Top motif match against the PLACE plant motif catalogue, determined using STAMP

§ STAMP *p*-value. Calculated from a test of the alignment between the input motif and the match

**Supplemental Table 2: List of putative wheat circadian oscillator genes.**

| Mace Gene ID | CS Gene ID | Gene Name |
|---|---|---|
| *TraesMAC6A03G03277000* | *TraesCS6A02G121500* | *TaADO-A1* |
| *TraesMAC6B03G03478600* | *TraesCS6B02G149800* | *TaADO-B1* |
| *TraesMAC6D03G03678160* | *TraesCS6D02G111600* | *TaADO-D1* |
| *TraesMAC7A03G03984110* | *TraesCS6B02G426300* | *TaADO-A2* |
| *TraesMAC7B03G04196410* | *TraesCS7A02G431600* | *TaADO-B2* |
| *TraesMAC7D03G04439440* | *TraesCS7D02G423400* | *TaADO-D2* |
| *TraesMAC6A03G03350200* | *TraesCS6A02G233800* | *TaCHE-A1* |
| *TraesMAC6B03G03553430* | *TraesCS6B02G262600* | *TaCHE-B1* |
| *TraesMAC6D03G03735860* | *TraesCS6D02G216100* | *TaCHE-D1* |
| *TraesMAC6A03G03414290* | *TraesCS6A02G412800* | *TaCHE-A2* |
| *TraesMAC6B03G03637890* | *TraesCS6B02G462100* | *TaCHE-B2* |
| *TraesMAC6D03G03797440* | *TraesCS6D02G397000* | *TaCHE-D2* |
| *TraesMAC1A03G00181960* | *TraesCS1A02G443200* | *TaELF3-A1* |
| *TraesMAC1B03G00403360* | *TraesCS1B02G477400* | *TaELF3-B1* |
| *TraesMAC1D03G00576650* | *TraesCS1D02G451200* | *TaELF3-D1* |
| *TraesMAC4A03G02066750* | | *TaELF4-L-A1* |
| *TraesMAC4B03G02286810* | *TraesCS4B02G149300* | *TaELF4-L-B1* |
| *TraesMAC4D03G02463080* | *TraesCS4D02G149000* | *TaELF4-L-D1* |
| *TraesMAC5A03G02664220* | *TraesCS5A02G201900* | *TaELF4-L-A2* |
| *TraesMAC5B03G02889420* | *TraesCS5B02G200500* | *TaELF4-L-B2* |
| *TraesMAC5D03G03116890* | *TraesCS5D02G208200* | *TaELF4-L-D2* |
| *TraesMAC4A03G02089120* | *TraesCS4A02G164000* | *TaFKF1-A1* |
| *TraesMAC4B03G02303040* | *TraesCS4B02G157500* | *TaFKF1-B1* |
| *TraesMAC3A03G01342510* | *TraesCS3A02G116300* | *TaGI-A1* |
| *TraesMAC3B03G01584430* | *TraesCS3B02G135400* | *TaGI-B1* |
| *TraesMAC3D03G01831120* | *TraesCS3D02G118200* | *TaGI-D1* |
| *TraesMAC5A03G02685150* | *TraesCS5A02G265500* | *TaJMJD5-A1* |
| *TraesMAC5B03G02915050* | *TraesCS5B02G265200* | *TaJMJD5-B1* |
| *TraesMAC5D03G03140140* | *TraesCS5D02G273400* | *TaJMJD5-D1* |
| *TraesMAC7A03G03928670* | *TraesCS7A02G299400* | *TaLHY-A1* |
| *TraesMAC7B03G04136500* | *TraesCS7B02G188000* | *TaLHY-B1* |
| *TraesMAC7D03G04386390* | *TraesCS7D02G295400* | *TaLHY-D1* |
| *TraesMAC4A03G02085650* | *TraesCS4A02G162200* | *TaLNK1-A1* |
| *TraesMAC4B03G02300160* | *TraesCS4B02G153700* | *TaLNK1-B1* |
| *TraesMAC4D03G02487410* | *TraesCS4D02G166000* | *TaLNK1-D1* |
| *TraesMAC3A03G01388770* | *TraesCS3A02G195700* | *TaLNK2-A1* |
| *TraesMAC3B03G01631750* | *TraesCS3B02G227900* | *TaLNK2-B1* |
| *TraesMAC3A03G01523810* | *TraesCS3A02G526600* | *TaLUX-A1* |

| | | |
|---|---|---|
| *TraesMAC3B03G01790490* | *TraesCS3B02G594300* | *TaLUX-B1* |
| *TraesMAC3D03G02002020* | *TraesCS3D02G531900* | *TaLUX-D1* |
| *TraesMAC1A03G00120120* | *TraesCS1A02G258400* | *TaLUX-L-A1* |
| *TraesMAC1B03G00320350* | *TraesCS1B02G268900* | *TaLUX-L-B1* |
| *TraesMAC1D03G00510100* | *TraesCS1D02G257700* | *TaLUX-L-D1* |
| *TraesMAC3A03G01460970* | *TraesCS3A02G361800* | *TaLUX-L-A2* |
| *TraesMAC3D03G01936510* | *TraesCS3D02G355500* | *TaLUX-L-D2* |
| *TraesMAC6A03G03317530* | *TraesCS6A02G193300* | *TaLWD-A1* |
| *TraesMAC6B03G03518210* | *TraesCS6B02G221400* | *TaLWD-B1* |
| *TraesMAC6D03G03715250* | | *TaLWD-D1* |
| *TraesMAC2A03G00601530* | *TraesCS2A02G081900* | *TaPpd-A1* |
| *TraesMAC2B03G00849230* | *TraesCSU02G196100* | *TaPpd-B1* |
| *TraesMAC2D03G01101810* | *TraesCS2D02G079600* | *TaPpd-D1* |
| *TraesMAC5A03G02704450* | *TraesCS5A02G320300* | *TaPRR59-A1* |
| *TraesMAC5B03G02934070* | *TraesCS5B02G320500* | *TaPRR59-B1* |
| *TraesMAC5D03G03158380* | *TraesCS5D02G326200* | *TaPRR59-D1* |
| *TraesMAC4A03G02039320* | *TraesCS4A02G105300* | *TaPRR73-A1* |
| *TraesMAC4B03G02333170* | *TraesCS4B02G198700* | *TaPRR73-B1* |
| *TraesMAC4D03G02504580* | *TraesCS4D02G199600* | *TaPRR73-D1* |
| *TraesMAC4A03G02110360* | *TraesCS4A02G200200* | *TaPRR95-A1* |
| *TraesMAC4B03G02269330* | *TraesCS4B02G115100* | *TaPRR95-B1* |
| *TraesMAC4D03G02447570* | *TraesCS4D02G112800* | *TaPRR95-D1* |
| *TraesMAC2B03G01023440* | *TraesCS2B02G478700* | *TaRVE27-B1* |
| *TraesMAC2D03G01259800* | *TraesCS2D02G456900* | *TaRVE27-D1* |
| *TraesMAC6A03G03361070* | *TraesCS6A02G261100* | *TaRVE27-A2* |
| *TraesMAC6B03G03565140* | *TraesCS6B02G288500* | *TaRVE27-B2* |
| *TraesMAC6D03G03746470* | *TraesCS6D02G241900* | *TaRVE27-D2* |
| *TraesMAC7A03G04029260* | *TraesCS7A02G553800* | *TaRVE27-A3* |
| *TraesMAC7B03G04256300* | *TraesCS7B02G478200* | *TaRVE27-B3* |
| *TraesMAC7D03G04483650* | *TraesCS7D02G540700* | *TaRVE27-D3* |
| *TraesMAC6A03G03360220* | *TraesCS6A02G258600* | *TaRVE68-A1* |
| *TraesMAC6B03G03564120* | *TraesCS6B02G266800* | *TaRVE68-B1* |
| *TraesMAC6D03G03745630* | *TraesCS6D02G239800* | *TaRVE68-D1* |
| *TraesMAC4A03G02215160* | *TraesCS4A02G474100* | *TaRVE86-A1* |
| *TraesMAC7A03G03805900* | *TraesCS7A02G017600* | *TaRVE86-A2* |
| *TraesMAC7D03G04266510* | *TraesCS7D02G014900* | *TaRVE86-D2* |
| *TraesMAC7A03G03999300* | *TraesCS7A02G470700* | *TaRVE86-A3* |
| *TraesMAC7B03G04214400* | *TraesCS7B01G372500* | *TaRVE86-B3* |
| *TraesMAC6A03G03345360* | *TraesCS6A02G227900* | *TaTOC1-A1* |
| *TraesMAC6B03G03549430* | *TraesCS6B02G253900* | *TaTOC1-B1* |

| | | |
|---|---|---|
| *TraesMAC6D03G03731950* | *TraesCS6D02G207100* | *TaTOC1-D1* |

**Supplemental Table 3: Allelic information for clock gene markers used to build chronotypes.** *TaPpd-B1*, *TaPpd-D1* and *TaELF3-D1* are existing major gene markers for wheat. *TaLUX-B1*, *TaPRR73-A1*, *TaTOC1-B1* and *TaPRR59-B1* are novel SNP markers identified from the OzWheat diversity panel.

| Allele | Mace gene ID | Chinese Spring gene ID | Variant type | Details | Heading phenotype | Reference |
|---|---|---|---|---|---|---|
| *TaPpd-B1* | *TraesMAC2B03G00849230* | *TraesCSU02G196100* | Copy number | | *a* & *c* head earlier than *b* & *d* | Cane *et al.* (2013) |
| *a* | | | | 3 copies | | |
| *b* | | | | 1 copy | | |
| *c* | | | | 4 copies | | |
| *d* | | | | 2 copies | | |
| *TaPpd-D1* | *TraesMAC2D03G01101810* | *TraesCS2D02G079600* | Indel | | *a* heads earlier than *b* and *d* | Cane *et al.* (2013); Beales *et al.* (2007) |
| *a* | | | | 2089 bp promoter deletion | | |
| *b* | | | | Intact | | |
| *d* | | | | 5 bp deletion in exon 7, generates a stop codon upstream of the CCT domain | | |
| *TaELF3-D1* | *TraesMAC1D03G00576650* | *TraesCS1D02G451200* | Inversion and indel | | *del* heads earlier than *WT* | Wittern *et al.* (2023) |
| *WT* | | | | Intact | | |
| *del* | | | | 2.6 Mbp inversion, containing a deletion in intron 2 | | |
| *TaLUX-B1* | *TraesMAC3B03G01790490* | *TraesCS3B02G594300* | SNP | | Unknown | |
| *T* | | | | Same as Refseq | | |
| *C* | | | | CDS change: T>C at pos. 247; Peptide change: S83P | | |
| *TaPRR73-A1* | *TraesMAC4A03G02039320* | *TraesCS4A02G105300* | SNP | | Unknown | |
| *G* | | | | Same as Refseq | | |
| *A* | | | | CDS change: G>A at pos. 1928; Peptide change: R643Q | | |
| *TaTOC1-B1* | *TraesMAC6B03G03549430* | *TraesCS6B02G253900* | SNP | | Unknown | |
| *A* | | | | Same as Refseq | | |
| *G* | | | | CDS change: A>G at pos. 1213; Peptide change: S405G | | |
| *TaPRR59-B1* | *TraesMAC5B03G02934070* | *TraesCS5B02G320500* | SNP | | Unknown | |
| *G* | | | | Same as Refseq | | |
| *A* | | | | CDS change: G>A at pos. 1297; Peptide change: G433S | | |

**Supplemental Table 4: Genotypes of the circadian diversity panel.** Genotypic information for the seven clock gene markers used to generate chronotypes. Details of clock gene variants are available in Supplemental Table 3.

| Cultivar | *Ppd-B1* | *Ppd-D1* | *ELF3-D1* | *LUX-B1* | *PRR73-A1* | *TOC1-B1* | *PRR59-B1* | Chronotype |
|---|---|---|---|---|---|---|---|---|
| Corack | *B* | *a* | *wt* | *T* | *G* | *A* | *A* | 1 |
| EGA Wedgetail | *b* | *a* | *wt* | *T* | *G* | *A* | *A* | 1 |
| Hydra | *b* | *a* | *wt* | *T* | *A* | *A* | *G* | 1 |
| Magenta | *b* | *a* | *wt* | *T* | *G* | *A* | *G* | 1 |
| Wyalkatchem | *b* | *a* | *wt* | *T* | *G* | *A* | *A* | 1 |
| Zen | *b* | *a* | *wt* | *T* | *G* | *A* | *A* | 1 |
| Scout | *b* | *a* | *del* | *T* | *A* | *A* | *A* | 2 |
| Yitpi | *b* | *d* | *del* | *C* | *A* | *A* | *A* | 2 |
| Buckley | *a* | *d* | *wt* | *C* | *A* | *G* | *G* | 3 |
| Insignia | *b* | *d* | *wt* | *T* | *A* | *G* | *G* | 3 |
| Viking | *b* | *d* | *wt* | *T* | *A* | *G* | *G* | 3 |
| Axe | *a* | *a* | *wt* | *C* | *G* | *A* | *A* | 4 |
| Bayonet | *a* | *a* | *wt* | *T* | *A* | *A* | *A* | 4 |
| Cascades | *a* | *a* | *wt* | *T* | *A* | *A* | *G* | 4 |
| Gladius | *a* | *d* | *wt* | *T* | *G* | *A* | *A* | 4 |
| Lillimur | *a* | *a* | *del* | *T* | *G* | *A* | *G* | 4 |
| Mace | *a* | *a* | *wt* | *T* | *G* | *A* | *A* | 4 |
| Sunguard | *a* | *a* | *wt* | *T* | *G* | *A* | *G* | 4 |
| Cobra | *a* | *b* | *del* | *T* | *G* | *A* | *A* | 5 |
| Takari | *b* | *a* | *del* | *T* | *G* | *G* | *G* | 5 |
| Wilgoyne | *a* | *a* | *del* | *T* | *A* | *A* | *A* | 5 |
| Calingiri | *d* | *a* | *wt* | *T* | *G* | *A* | *G* | 6 |
| Suntop | *d* | *a* | *wt* | *T* | *G* | *A* | *G* | 6 |
| Supreme | *d* | *a* | *wt* | *T* | *A* | *A* | *G* | 6 |
| Yarralinka | *c* | *d* | *wt* | *T* | *G* | *A* | *G* | 6 |

**Supplemental Table 5: Genotypes of major clock and phenology genes for each NIL.** Allelic information for clock gene markers is available in Supplemental Table 3.

| Line | Pair | Cross | *Ppd-B1* | *Ppd-D1* | *VRN-A1* | *VRN-B1* | *VRN-D1* | *ELF3-D1* |
|---|---|---|---|---|---|---|---|---|
| CSIROW173 | a | 17x7374x28709 | *b* | *b* | *v* | *v* | *a* | *WT* |
| CSIROW084 | | 17x7374 | *b* | *b* | *v* | *v* | *a* | *del* |
| CSIROW185 | b | 17x7374x28709 | *b* | *b* | *v* | *v* | *v* | *WT* |
| CSIROW046 | | 17x7374 | *b* | *b* | *v* | *v* | *v* | *del* |
| CSIROW174 | c | 17x7374x28709 | *b* | *a* | *v* | *a* | *a* | *WT* |
| CSIROW017 | | 7374 | *b* | *a* | *v* | *a* | *a* | *del* |
| CSIROW175 | d | 17x7374x28709 | *b* | *b* | *v* | *a* | *a* | *WT* |
| CSIROW075 | | 7374 | *b* | *b* | *v* | *a* | *a* | *del* |

**Supplemental Table 6: Novel primers developed for KASP genotyping.** Homoeologue-specific primers were generated using PolyMarker(Ramirez-Gonzalez et al., 2015).

| Name | Sequence | SNP name | Strand |
|---|---|---|---|
| LUX-B1_A1 | GGGGAAGGAGGGGAAGGAC | | |
| LUX-B1_A2 | GAAGGTGACCAAGTTCATGCTCGCTCCGCCTCCTCGT | CSIROSNP_16129 | Forward |
| LUX-B1_R | GAAGGTCGGAGTCAACGGATTCGCTCCGCCTCCTCGC | | |
| PRR73-A1_A1 | GGGTCGTGTCATCTGTCTG | | |
| PRR73-A1_A2 | GAAGGTGACCAAGTTCATGCTGAGACTTGCCGAGCAGCG | CSIROSNP_18574 | Forward |
| PRR73-A1_R | GAAGGTCGGAGTCAACGGATTGAGACTTGCCGAGCAGCA | | |
| PRR59-B1_A1 | TATCCACAGTCAGGTCCTCCTATC | | |
| PRR59-B1_A2 | GAAGGTGACCAAGTTCATGCTTGCGACACTGGAGTTCTGCC | CSIROSNP_24473 | Reverse |
| PRR59-B1_R | GAAGGTCGGAGTCAACGGATTTGCGACACTGGAGTTCTGCT | | |
| TOC1-B1_A1 | TCAAGCTCATACACCACCAAC | | |
| TOC1-B1_A2 | GAAGGTGACCAAGTTCATGCTCACATTCATACCAGCAGGACT | CSIROSNP_30035 | Reverse |
| TOC1-B1_R | GAAGGTCGGAGTCAACGGATTCACATTCATACCAGCAGGACC | | |

### Supplemental Methods

#### Full details of growth conditions

*Cultivar panels*

In year one, plants were grown in a glasshouse (BioSciences 3, University of Melbourne, Parkville; 37°47'49.6"S 144°57'36.1"E). Seeds were sown in two batches of 75 on 10th July 2020 and 17th July 2020. Plants were grown in 500 mL pots arranged according to a randomised complete block design (RCBD). Each pot received 15 mL tap water twice daily via an automated irrigation system. Typical light intensity across the tables at midday ranged between 180-220 µmol $m^{-2}$ $s^{-1}$. Circadian rhythms were measured in four replicates of each cultivar from each batch.

In years two and three, plants were grown in a Glasshouse (BioSciences 4, University of Melbourne, Parkville; 37°47'49.4"S 144°57'32.0"E). In the second year, circadian rhythm, senescence, nutrient mobilisation and GPC phenotyping was performed. In the third year, circadian rhythm phenotyping was performed. In year two, seeds were sown in four batches on 15th July 2021, 29th July 2021, 5th August 2021 and 12th August 2021. The first batch comprised 12 replicates of each cultivar, sown in 1 L pots, which were used for senescence and nutrient profiling experiments. The second, third and fourth batches each comprised four replicates of each cultivar, sown in 500 mL pots, which were used for circadian rhythm phenotyping experiments. All pots were arranged according to a RBCD. Each pot received 30 mL tap water (1 L pots) or 15 mL tap water (500 mL pots) twice daily via an automated irrigation system. Supplementary lighting was used between the hours of 8.30 AM and 4.30 PM (with lights on after dawn and lights off prior to dusk) throughout the growing year. Typical light intensity across the tables at midday ranged between 280-350 µmol $m^{-2}$ $s^{-1}$.

In year three, seeds were sown in two batches on 29th July 2022 and 5th August 2022. Each batch comprised five replicates of each cultivar, sown in 1 L pots arranged according to a RBCD. Each pot received 30 mL tap water twice daily via an automated irrigation system. Typical light intensity at midday was 300-350 µmol $m^{-2}$ $s^{-1}$. Plants were positioned in a brighter area of the glasshouse in year three and supplementary lighting was not required.

In year four, plants were grown in a controlled environment room. Senescence, GPC and TGW phenotyping was performed. Room settings were 12 h light (20°C): 12 h dark (15°C) cycles, and light intensity was 200-280 µmol $m^{-2}$ $s^{-1}$. Five replicates of each cultivar were sown in 1 L pots. Plants received approximately 30 mL tap water twice daily via an automated irrigation system.

*Developmental time course experiment*

For the developmental time course experiment, seed from three cultivars (Cobra, Mace and Supreme) was sown into 1 L pots (15 replicates per cultivar) in a controlled environment room (12 h light (20°C): 12 h dark (15°C) cycles, light intensity was 200-280 µmol $m^{-2}$ $s^{-1}$). Plants were divided into three batches of five replicates that were phenotyped at discrete developmental stages (described below).

*TaELF3-D1 NILs*

Five replicates of each of the eight NILs were sown into 1 L pots in a Conviron controlled environment cabinet. Light and temperature conditions were set to 12 h light (20°C): 12 h dark (15°C) cycles, and light intensity ranged between 190-220 µmol $m^{-2}$ $s^{-1}$ across the cabinet. Pots were placed in trays (six pots per tray), and approximately 1 L tap water was applied to each tray three times weekly. Following circadian rhythm phenotyping, plants were relocated to a controlled environment room (12 h light (20°C): 12 h dark (15°C) cycles, light intensity was 200-280 µmol $m^{-2}$ $s^{-1}$) prior to senescence, GPC and TGW phenotyping.

*Circadian transcriptomic experiment*

Wheat seeds of the Australian wheat cultivar Mace were sown in two batches of 96 pots in a controlled temperature growth room. Batches were sown 28 days apart. The temperature and light conditions were set to 12 h light (20°C): 12 h dark (15°C) cycles, with light intensity ranging from 200-280 µmol $m^{-2}$ $s^{-1}$ across the growing area. Plants received approximately 30 mL water twice daily via an automated irrigation system.
